# Impacts of *Mycoplasma agalactiae* restriction-modification systems on pan-epigenome dynamics and genome plasticity

**DOI:** 10.1101/2021.06.21.448925

**Authors:** Emilie Dordet-Frisoni, Céline Vandecasteele, Rachel Contarin, Eveline Sagné, Eric Baranowski, Christophe Klopp, Laurent Xavier Nouvel, Christine Citti

## Abstract

DNA methylation plays an important role in the biology of bacteria. Often associated with restriction modification (RM) systems, they also provide a defence against foreign DNA. Little is known regarding the methylome of the mycoplasma genus, which encompasses several pathogenic species with small genomes. Here, single molecule real-time (SMRT) and bisulphite sequencing combined with whole-genome analysis identified 19 methylated motifs associated with three orphan methyltransferases (MTases) and eight RM systems in *Mycoplasma agalactiae*, a ruminant pathogen and a model organism. All systems had a homolog in at least one phylogenetically distinct *Mycoplasma* spp. Our study also revealed that several superimposed genetic events may participate in the *M. agalactiae* dynamic epigenome landscape. These included (i) DNA shuffling and frameshift mutations that affect the MTase and restriction endonuclease content of a clonal population and (ii) gene duplication, erosion, and horizontal transfer that modulate MTase and RM repertoires of the species. Some of these systems were experimentally shown to play a major role in mycoplasma conjugative, horizontal DNA transfer. While the versatility of DNA methylation may contribute to regulating essential biological functions at cell and population levels, RM systems may be key in mycoplasma genome evolution and adaptation by controlling horizontal gene transfers.

## INTRODUCTION

*Mycoplasma* spp. represent a large group of bacteria that have adopted a parasitic lifestyle and closely interact with their human or animal hosts, as commensals or pathogens (1). These organisms belong to the Mollicutes class and encompass some of the simplest life forms capable of self-replicating under laboratory conditions in axenic media. From a phylogenetic point of view, mycoplasmas derived from a common ancestor to Gram-positive bacteria with a low GC content and followed an evolutionary scenario often described as a ‘degenerative evolution’ because of successive and drastic genetic losses. As a result, current mycoplasmas are characterised by a small genome (ca. 0.5 to 1.35 Mbp), no cell wall or cell wall precursors, and a restricted number of metabolic pathways (1). For decades, genome reduction has been proposed as the only force driving mycoplasma evolution. Horizontal gene transfer (HGT) was then considered marginal in these organisms because of their limited content in efficient recombination systems (2) and their paucity in mobile genetic elements (MGEs; 3, 4). A paradigm shift occurred within the last 20 years with the first discoveries of conjugative MGEs in mycoplasmas (5, 6) and large chromosomal DNA exchanges between phylogenetically remote *Mycoplasma* spp. sharing the same habitat (4, 7, 8). *In vitro* experiments further demonstrated that mycoplasmas harbouring integrative and conjugative elements (ICEs) have retained a form of sexual competence (9, 10). Most specifically, our group highlighted two conjugative processes occurring within and among strains of *M. agalactiae*, an important pathogen of ruminants and a model organism. The first was the conventional, horizontal dissemination of mycoplasma ICEs (MICEs), from ICE-positive to ICE-negative mycoplasma cells (9, 11). The second involved the transfer of chromosomal DNA (10, 12), a conjugative process initially described from ICE-negative to ICE-positive cells and further designated as MCT for mycoplasma chromosomal transfer (13). MCT is an atypical mechanism of HGT that is not physically linked to MGE movements but relies on the MICE conjugative machinery. For MCT to occur, one mating partner must carry a functional MICE (9, 11). The mechanism driving MCT remains to be fully understood, but its outcome has been well described. Multiple, large, and small chromosomal regions of the recipient cell are simultaneously replaced by the donor-counterparts via recombination, as described for the distributive conjugative transfer (DCT) in Mycobacteria (14, 15). While the frequency of such a phenomenon is low, the impact of MCT has far-reaching consequences, as the mating of two strains is able to generate progenies composed of a variety of individual mosaic genomes, each being unique (12, 13). One recurrent observation was the apparent polarity of the chromosomal transfer during mating experiments involving *M. agalactiae* strain 5632, with this particular strain always being identified as the recipient genome regardless of its mating partner. The same was observed when conjugation was bypassed by PEG-cellular fusion, suggesting that the asymmetry of the chromosomal transfer might be independent of the conjugative mechanism itself (13).

Restriction modification (RM) systems are key players of bacterial HGT by protecting their host genome from incoming MGEs or by promoting recombination of the invading DNA, as recently suggested (16, 17). Furthermore, detailed analyses suggest that the small genome of strain 5632 has a relatively large arsenal of RM systems (18), raising the question of their role in the apparent polarity of MCTs.

RM systems are classically composed of two elements: a restriction endonuclease (REase) and a methyltransferase (MTase), which both usually bind to the same DNA sequence. They are classified into four major types (Type-I to -IV RM systems), according to their subunit composition, sequence recognition strategy, substrate specificity, and cleavage position (19). More specifically, MTases induce three types of base methylations: *N*6-methyladenine (6mA), C5-methyl-cytosine (5mC), and *N*4-methyl-cytosine (4mC), the latter being only found in bacteria and archaea. Most of these methylated bases are induced by DNA adenine MTase (Dam) or DNA cytosine methyltransferase (Dcm) encoded by specific *dam* and *dcm* genes. Besides protecting genomic DNA from cleavage and degradation by cognate REases, MTases also contribute to several bacterial processes, such as DNA mismatch repair, gene regulation, replication initiation, cell cycle progression, and phase variation (20). The small mycoplasma genome lacks most known transcription factors and regulatory pathways. Despite this limitation, mycoplasmas can respond to environmental stress and metabolic insults (21–23), suggesting other types of regulation, such as DNA methylation, as epigenetic modulators of gene expression.

Based on comparative genomic analyses, the Tenericutes phylum (class Mollicutes) includes the Mycoplasma genus and has a high density of RM systems, a puzzling observation regarding the reduced genome sizes of these peculiar bacteria (16). In this phylum, RM systems are ubiquitous and found in 74.2% of the genomes. Their distribution patterns are very diverse, with more than 2,040 MTases in the Restriction Enzyme database (REBASE) that were predicted from 387 sequenced mycoplasma genomes (24), and an average of five predicted MTases per genome. While this highlights the importance of MTtases and RM systems in mycoplasmas, their role in cellular regulation and HGT has yet to be addressed. Genome methylation in bacteria is an area of great interest because of its broad implications for evolution, biology, and virulence (25). The development of real-time single molecule sequencing (SMRT-seq) has allowed the detection of methylated bases, whether on a plasmid or bacterial chromosome, and is particularly suited to small mycoplasma genomes. However, the characterisation of Mycoplasma epigenomes at a single-base resolution is still scarce, with only two comprehensive studies. The first one was conducted in 2013 with the type strains of two human *Mycoplasma* spp. (26), *M. genitalium* and *M. pneumoniae*. This study revealed two different m6A methylated motifs, one common to both bacteria (5’-CTA^m6^T-3’) and one specific to *M. pneumoniae* that corresponded to the Type-I methylated motif (5’-CT(N)_7_A^m6^TR-3’). Functional and distribution analyses further suggested a potential regulatory role for these methylations in cell cycle and gene expression. More recently, the *M. bovis* reference strain PG45 was included in a large-scale epigenetic analysis comprising more than 200 bacterial and archaeal genomes, which were analysed by SMRT-seq (27); five different methylated motifs, four with m6A and one with m4C, were identified for 14 putative MTases identified in the PG45 genome. While providing the first insights into the mycoplasma epigenome, these studies were limited to Pacbio SMRT-seq, which does not allow the detection of m5C modifications, and to a single strain per species.

The current study provides new insights into the extent, variability, and impacts on gene transfers of DNA methylation in *M. agalactiae*, a model species for the understanding of mycoplasma HGTs. This was achieved by combining SMRT-seq and Illumina bisulphite sequencing (BS-seq) for the detection of all cytosine modifications (m4C and m5C) in addition to m6A. In-depth analysis of the *M. agalactiae* epigenome landscape was conducted using two reference strains, namely 5632 and PG2, for which circularised genomes were available, as well as data on HGT and genetic tools. Eight additional field strains, with varied histories, were also used to assess the methylome diversity within the *M. agalactiae* species and identify the repertoire of active RM systems along their recognition motifs. This knowledge was then used to address the impact of RM systems on HGTs by experimentally demonstrating the influence of DNA methylation on the outcome of mycoplasma conjugation and by establishing a correlation between mobilomes and active RM repertoires of *M. agalactiae* genomes. MTases identified in this study all have homologs in at least one other *Mycoplasma* spp. of a different phylogenetic group, indicating that data obtained with *M. agalactiae* can be extrapolated to several other Mollicutes. Altogether, our data highlight the intra-clonal and intra-species versatility of the *M. agalactiae* epigenome and unveil the dual role played by RM systems in HGT. While some RM systems provide the mycoplasma with an ‘immune system’ to prevent foreign DNA from entering, others may play a major role in HGT by promoting the incorporation of the donor chromosomal DNA into the host genome.

## MATERIALS AND METHODS

### Mycoplasma strains, culture conditions, and DNA extraction

The *Mycoplasma agalactiae* strains used in this study are described in Table 1. The 5632 H1-2 (13) and the PG2-ICEA+ (9) variants were previously generated and derived from strain 5632 and PG2, respectively. Briefly, the 5632 H1-2 variant is a hybrid obtained by PEG-cellular fusion of 5632 with PG2 and is characterised by the 5632 genomic-background in which the *hsd* locus has been replaced by its PG2 counterpart, in addition to four other unrelated regions (13). PG2-ICEA+ is a PG2 variant having acquired and integrated a functional ICEA_5632_ after conjugation with 5632. All strains and variants were grown for 24 to 48 h at 37°C in SP4 medium supplemented with cephalexin (500 µg.ml^-1^). When necessary, strains were subcloned by serial passages in broth and solid media. All mycoplasma cultures were stored at −80°C.

**Table 1.** Features of *Mycoplasma agalactiae* strains used in this study

| Strains | Host | Tissue sample<br>(Associated symptoms) | Date of isolation | Geographical origin ( | ST | Estimated genome size (bp) | Number of contigs | %GC | N50 value | Number of CDS |
| --- | --- | --- | --- | --- | --- | --- | --- | --- | --- | --- |
| **5632** | Caprine | Joint | 1991 | Spain | ST-13 | 1006702 | 1<br>circularised | 29.7 |  | 752 |
| **PG2** | Caprine | Unknown | 1952 | Spain | ST-06 | 877438 | 1<br>circularised | 29.62 |  | 826 |
| **4021** | Ovine | Milk<br>(asymptomatic) | 1988 | France (12) | ST-10 | 887230 | 3 | 29.7 | 822961 | 846 |
| **4025** | Ovine | Milk<br>(asymptomatic) | 1988 | France (12) | ST-11 | 887230 | 3 | 29.7 | 822961 | 984 |
| **4055** | Caprine | Milk (mastitis) | 1987 | France | ST-03 | 926904 | 1 | 28.6 | 926904 | 3017 |
| **4210** | Ovine | Milk (mastitis) | 1982 | France (73) | ST-05 | 884408 | 4 | 29.7 | 614215 | 819 |
| **5276** | Ovine | Milk (mastitis) | 1977 | France | ST-12 | 890157 | 3 | 29.7 | 781215 | 823 |
| **13377** | Caprine | Ear | 2003 | France (79) | ST-08 | 1266155 | 20 | 29.6 | 115104 | 1203 |
| **14634** | Ovine | Milk (mastitis) | 2006 | France (64) | ST-12 | 905745 | 6 | 29.7 | 615197 | 838 |
| **14668** | Caprine | Lung<br>(pneumonia) | 2006 | France (7) | ST-14 | 967428 | 4 | 29.8 | 808260 | 912 |

ST : Shared type determined for European *M. agalactiae* strains in Nouvel *et al.* (66)
CDS : Coding DNA sequence
N50 value : shortest contig length needed to cover 50% of the genome.

Genomic DNA was extracted from mycoplasma cells after 48 h of growth (beginning of stationary phase) using the phenol-chloroform method as previously described (29). To assess the methylation status, depending on the growth phase of *M. agalactiae*, chromosomal DNA of 5632 was also extracted after 24 h, during the exponential growth phase.

### SMRT, BS-seq, and sequence annotation

Genome sequencing of *M. agalactiae* strains was performed at the GeT-PlaGe Genotoul facility (Toulouse) using Pacific Biosciences SMRT RSII technology. The SMRTbell library was constructed by pooling 20 barcoded samples (10 were used in this study). Barcoded library preparation and sequencing were performed according to the manufacturer’s instructions “Preparing SMRTbell(tm) Libraries using PacBio^®^ Barcoded Adapters for Multiplex SMRT^®^ Sequencing.” Genomic DNA was quantified at each step using the Qubit dsDNA HS Assay Kit (LifeTechnologies, Courtaboeuf, France) and a Qubit Fluorometer (ThermoFischer Scientific). DNA purity was tested using a Nanodrop (Thermofisher). Size distribution and degradation were assessed using a fragment analyser (AATI) High Sensitivity NGS Fragment Analysis Kit. The DNA of each sample was sheared at 9 kb using the Megaruptor system (Diagenode) and then barcoded using the SMRTbell Barcoded Adapter Prep Kit (Pacific Bioscience, Menlo Park, USA). Briefly, each sample underwent one-step end repair, and the barcode adapters were ligated. The barcoded samples were then pooled in equal amounts to obtain a final pool, which was purified with AMPure PB beads at ×0.45 and quantified. This pool then underwent DNA damage repair and exonuclease digestion and was purified with AMPure PB beads at ×0.45. A size selection step using a 6 kb cut-off was performed on the Blue Pippin Size Selection system (Sage Science) with the 0.75% agarose cassettes, using Marker S1 high Pass 6-10 kb V3. The sized pool was purified again with AMPure PB beads at ×0.45 and quantified. Conditioned Sequencing Primer V2 was annealed to the size selected SMRTCell library. The annealed library was then loaded on four SMRTCells using P6–C4 chemistry on the PacBio RSII sequencer with 360-minute movies. Sequencing reads were *de novo* assembled using the Hierarchical Genome Assembly Process (HGAP) protocol RS_Assembly v.3 implemented in SMRT Analysis software v.2.3 with default parameters (https://github.com/PacificBiosciences/SMRT-Analysis). An average of 1 × 10^5^ reads per strain were obtained, corresponding to an average of 100X coverage.

Open reading frame (ORF) prediction and automatic annotation were performed using Rapid Annotation using Subsystem Technology (RAST; 30). GenBank editing and manual expert inspection were performed using Artemis 16.0.0 (31) and ACT 13.0.0 (32).

### Methylome analysis (SMRT and BS sequencing)

Methylome analysis was performed using a combination of Pacbio SMRT-seq, BS-seq, and comparative genome analysis. With the SMRT data, epigenetic modification at each nucleotide position was determined based on the kinetic variations (KVs) in the nucleotide incorporation rates, and methylated motifs were deduced from the KV data. SMRT-seq only identified m6A and m4C modifications. Base modifications and detection of methylated motifs were performed with SMRT analysis software (SMRT-v2.3.0) following *de novo* genome assembly using the ‘RS_Modification_and_Motif_Analysis.1’ protocol using a QV threshold of 60. A QV of 30 was also tested, but a QV of 60 was retained because it allowed the detection of N4-methylcytosine (m4C) in addition to N6-methyladenine (m6A). SMRT methylation motifs with a mean score of up to 40 (corresponding to a P-value of 0.0001) were considered specific and were used for further analysis (https://github.com/PacificBiosciences/Bioinformatics-Training/wiki/Methylome-Analysis-Technical-Note). The SMRT^®^View browser (v2.3.0) was used for visualisation and analysis of SMRT-detected methylated motifs. Notably, a bias was observed at each end of the contigs, with several sites not being detected as methylated within the first 2 and last 2 kbps. This was due to an intrinsic characteristic of SMRT-seq, whose coverage across a genome closely follows a Poisson distribution, resulting in a coverage decrease at each end of the contig. When the coverage in these areas was less than 25X per strand, the detection of methylated sites was not possible. This was, however, circumvented by comparing the 5632 and PG2 methylomes. Counterparts of contig extremities in one genome were 100% methylated in the other and vice versa, confirming the technical detection bias. This was possible because the organisation and sequence of the two genomes were very similar.

The 5632 methylome was evaluated after 24 or 48 h of growth to define the impact of the growth phase on the methylation status. SMRT analysis detected the same methylated motifs in the exponential phase (24 h) and the stationary phase (48 h), with a decrease of approximately 20% of the m6A methylated sites at 24 h and 40% for m4C (data not shown).

Illumina sequencing of bisulphite-treated chromosomal DNA (termed BS-seq) was performed to reliably detect 5-methylcytosine (m5C). Two reference strains 5632 and PG2 were used for BS-seq, and the 5632-like strains 13377, 4025, 14668, and 4055 were chosen based on dcm gene detection by BLAST analyses against the Type II m5C methyltransferase genes databases (dcm database) available on REBASE (Table S1; 24). BS-seq was performed at Novogene (UK) and included DNA fragmentation into 200–400 bp using Covaris S220, DNA libraries, and bisulphite treatments (EZ DNA Methylation Gold Kit, Zymo Research). Libraries were sequenced using a HiSeq 2500 platform, which enables paired-end sequencing and generates 125 bp long reads. The BS-seq raw data sets were analysed with Bismark software (v0.22.1) to detect m5C modifications and identify corresponding methylation motifs. Raw reads were mapped on the finished reference genomes using Bismark Bisulfite mapper v0.22.1 (https://www.bioinformatics.babraham.ac.uk/projects/bismark/) in non-directional mode. The methylated cytosines were extracted with bismark_methylation_extractor using --comprehensive, --merge_non_CpG, and --CX options. BedGraph and cytosine_report outputs were generated. The sequence context was analysed by extracting ± 10 bp around each identified methylated cytosine. The MEME-ChIP suite (Motif Analysis of Large Nucleotide Datasets, v5.1.0; http://meme-suite.org/) was used to determine a context consensus at methylated positions, and Artemis 16.0.0 (31) was used for visual inspection.

RM system-encoding genes and methylation motif(s) co-occurrence were addressed by comparative genome analysis. All identified ORFs were searched for similarity to known RM systems using BLASTP and the 5632 and PG2 reference genomes and the REBASE database (http://rebase.neb.com/rebase/rebase.html; 24). Significant BLASTP hits were selected using a cut-off E-value of <0.001 and exhibiting over 30% similarity across at least 80% of the sequence length. The active MTases were also analysed by BLASTP on NCBI (https://blast.ncbi.nlm.nih.gov) to determine their occurrence within Mollicutes and other bacterial classes. Significant BLASTP hits were selected using a cut-off E-value of <0.001 and exhibiting over 50% similarity across at least 50% of the sequence length.

### Sanger sequencing to validate the number of repeats in the poly(GA) or poly(G) regions

During *de novo* genome assembly, repetitive sequences can lead to erroneous rearrangements, deletions, and collapsed repeats, even with long read sequencing data. To validate the number of poly(GA) and poly(G) tracts detected in Type III and CpG MTases sequences, respectively, PCR was performed with primers described in Table S2 according to the Phusion^®^ High-Fidelity DNA polymerase supplier recommendations (New England Biolabs). Genomic DNA sequencing was performed using the 1F_typeIII primers (Table S2) by Eurofins genomics (Ebersberg, Germany).

### Transformation and complementation of *M. agalactiae* strains

Six MTase genes of 5632 (MAGa1570, MAGa1580, MAGa2700, MAGa3950, MAGa4250, and CDSH) were independently cloned into PG2, a strain deprived of these genes or having no active form (Table 2). As previously described, each of the six genes was placed under the control of the constitutive *M. agalactiae* P40 promoter (28) and inserted at the *Not*I restriction site of a mini-transposon. This mini-transposon is derived from the Tn4001 transposon, contains a gentamicin resistant gene (*aap*), and is located in the pMT85Gm, a plasmid that cannot replicate in mycoplasmas (33). Thus, complementation was obtained by inserting the mini-transposon and the MTase of interest in the genome. This construction has the advantage of (i) being stable because the transposase encoded by the non-replicative pMT85Gm is placed outside the mini-transposon and is lost after integration of the mini-transposon and (ii) preventing potential lethal effects due to enzyme overexpression when carried on a replicative plasmid.

**Table 2.**
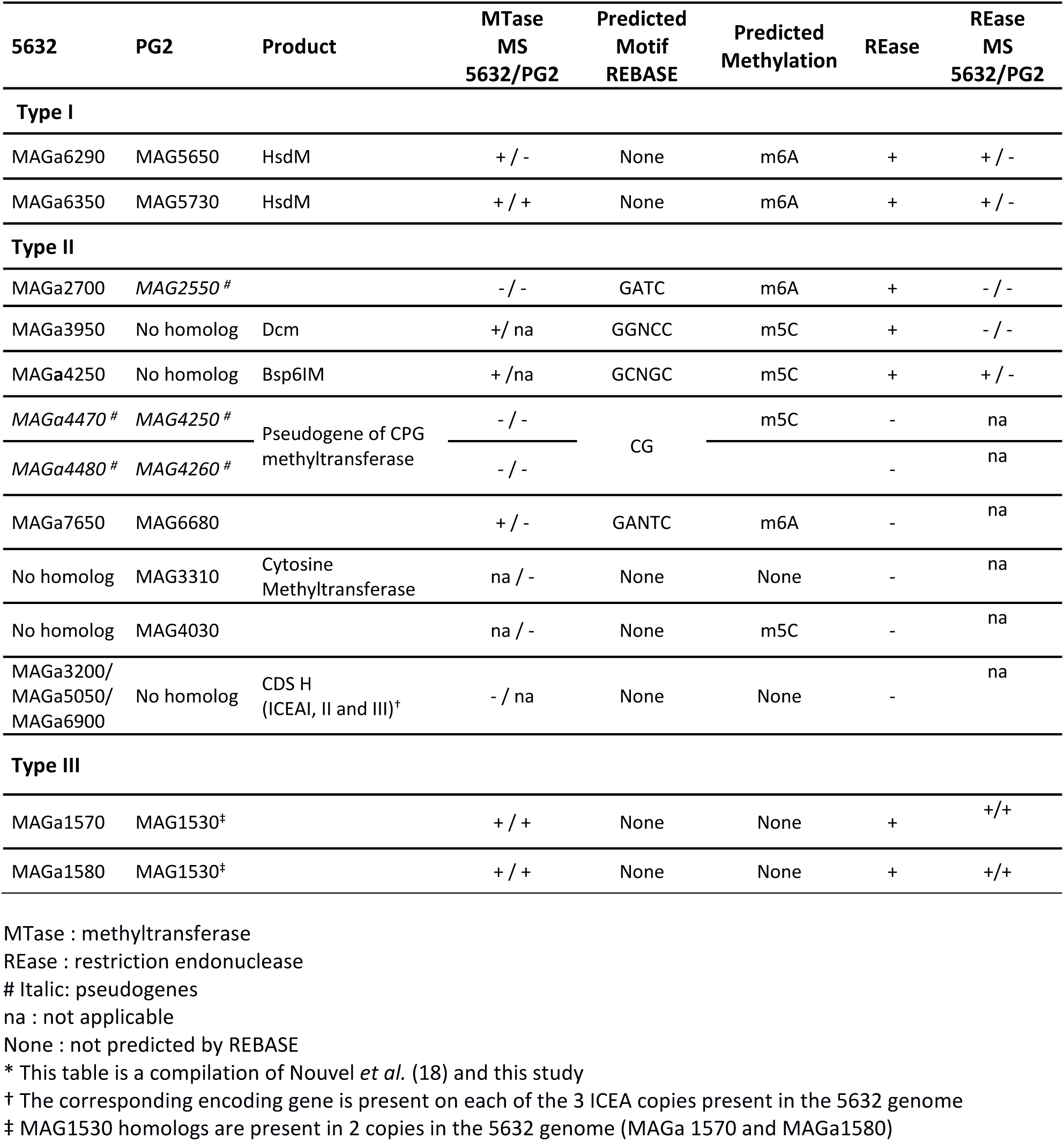
DNA methyltransferases detected by REBASE database in 5632 and PG2 genomes*

*M. agalactiae* transformation was performed as previously described, with some minor modifications (34). Briefly, a culture containing 10^8^ CFU/mL was centrifuged at 10,000 g at 4°C for 20 min. The pellet was washed twice with sterile cold DPBS 1X and centrifuged at 10,000 g at 4°C for 10 min. After centrifugation, cells were resuspended in 375 μL of cold 0.1 M CaCl_2_ and incubated on ice for 30 min. Cold CaCl_2_-incubated cells (100 μL) were gently mixed with 10 μg of yeast tRNA (Life Technologies, Carlsbad, CA, United States) and 3 μg of pMT85Gm MTase gene-harbouring plasmids or the pMT85Pur plasmid, which carry a puromycin resistance *pac* gene (GenBank: ACR82772.1). One mL of 50% PEG 8000 (Sigma-Aldrich, Saint-Louis, MO, United States) was then applied for 1 min, and the reaction was stopped by the addition of 5 mL SP4 liquid medium. The cells were incubated for 3 h at 37°C, and the transformation mix was centrifuged at 8,000 g at room temperature for 8 min. The pellet was suspended in 1 mL SP4 liquid medium and 300 μL were plated onto selective SP4 solid medium supplemented with 50 µg/mL gentamicin or 5 μg/mL puromycin (Sigma-Aldrich, Saint-Louis, MO, United States) and incubated at 37°C for 3 to 5 days. Colonies obtained on selective solid media were picked and transferred into 1 mL SP4 liquid medium supplemented with gentamicin (50 µg.mL^-1^), and incubated at 37°C for 24 to 96 h. Transformants were stored at −80°C.

To demonstrate the association between MTases and cognate methylated motifs and to validate the complementation efficiency, restriction assays were conducted in parallel. For this purpose, chromosomal DNA, extracted from PG2 and complemented PG2 clones using the phenol-chloroform method (29), was subjected to commercially available enzymes, *Sau*96I, *Fnu4*HI, *Dpn*I, and *Dpn*II according to the supplier’s recommendations (New England Biolabs) and analysed by agarose gel electrophoresis 1% TAE 0.5X (Figure S1). For the other MTases, there was no available commercial enzyme that corresponded to the recognition site, and for one, CDSH, the corresponding methylation site was not known.

### Conjugation experiments with PG2 clones complemented with 5632 MTases

Mating experiments were conducted as previously described (35) using the 5632TH3 tetracycline-resistant clone (13) as the recipient strain and, for each complemented MTase, a pool of ten PG2 clones carrying a single copy of the 5632 MTase gene inserted into the chromosome. This strategy was used to avoid potential negative effects linked to mini-transposon insertion that occurred at random and might affect growth or conjugation. The mating efficiency was determined as the number of transconjugants divided by the total CFUs and compared to a control performed with 5632TH3 and a pool of ten PG2 clones integrated with the non-complemented Gm^R^ transposon at different chromosomal positions. The results were expressed as a percentage (5632TH3 x PG2+MTase versus 5632TH3 x PG2 control), with 100% representing no difference; the lower the percentage, the greater the effect of MTase mycoplasma chromosomal transfer.

### Correlation analyses between HGT and RM systems in *M. agalactiae*

Mobile elements present in *M. agalactiae* genomes were identified after annotation using the RAST server (30). CDS annotated ‘mobile elements’ by RAST but also integrases, transposases, and prophage elements were counted, as well as genes identified as ICE by BLAST analyses against 5632 ICEA (e-value superior to 10^−3^). ICEs were considered when at least three of their essential components that are conserved across mycoplasma ICEs (CD5, CDS17, and CDS22) were present within a 30 kb DNA fragment. These were CDS5 and CDS17 encoding the conjugation factors TraE and TraG and CDS22 that encodes the DDE transposase (11). Correlation analyses were performed with GraphPad Prism version 6.00 (GraphPad Software, San Diego, California, USA).

## RESULTS AND DISCUSSION

### *M. agalactiae* genome-wide methylation landscape

A previous comparative genomic analysis conducted by our group suggested that *M. agalactiae* strains varied in their repertoire of RM systems (18), a situation that may impact their epigenomes differently. To gain insight into the epigenome of this species, DNA methylation profiles were first defined with two representative strains for which complete, circularised genomes were already available, namely 5632 and PG2. For this purpose, a combination of Pacbio SMRT-seq and Illumina BS-seq, two complementary approaches, were used. SMRT-seq identified m6A and m4C modifications, while BS-seq detected both m4C and m5C. Intra-species diversity of the *M. agalactiae* pan-epigenome was further investigated by extending this approach to eight additional strains having different histories and collected over 30 years in France (Table 1).

Methylome data first indicated that the 5632 genome has twice as many methylated sites when compared to PG2, with a total of 26,444 and 13,268 sites detected, respectively. This observation correlated with 5632 having a higher number of predicted MTases than PG2 (18; Table 2). Combined with the sequence context, DNA methylations were associated with 13 unique motifs in both strains (Table 3), six of which were assigned to the Type I RM system and seven to Types II and III. For both strains, the classical m6A adenine modification (dam methylation) was the most prevalent, as observed for other bacteria (27). In contrast, cytosine modifications (dcm methylation) were detected in 5632 but not in PG2 and included the m4C modification identified by both SMRT- and BS-seq in the 5’-GC^m4^NGC-3’ motif, and the m5C modification was only detected by BS-seq in the 5’-GGNC^m5^C-3’ (Table 3). Regardless of the methylation type, perfect palindromes were found to be methylated on both strands (dimethylated), while imperfect palindromes such as 3’-RCA^m6^C-5’and 3’-GA^m6^AG-5’ were hemimethylated (Table 4). A mix of both hemi- and dimethylated sites was observed for some Type II motifs, such as 5’-YRA^m6^TC-3’ and Type I 5’-HA^m6^YC(N)_5_KTAA-3’. Whether this situation reflects a difference in MTase affinity for their recognition sequence when more than one base is permitted at a particular position of the target site (degenerated recognition sequences) or differences in DNA methylation of both strands during DNA replication is not known.

**Table 3.**
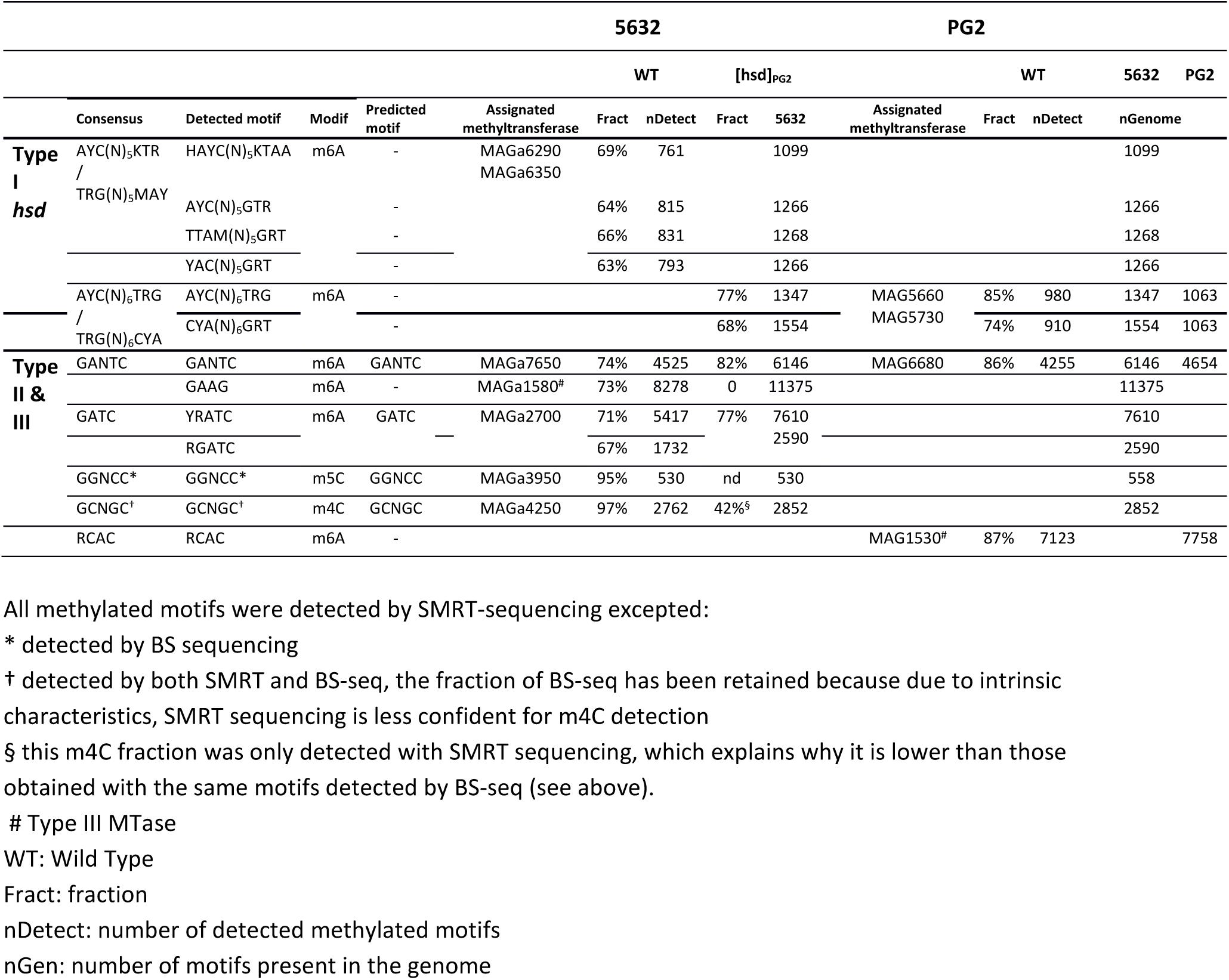
Methylated motifs detected in *M. agalactiae* 5632, 5632[hsd]_PG2_ (variant H1-2), and PG2 strains by SMRT and Bisulphite sequencing

**Table 4.**
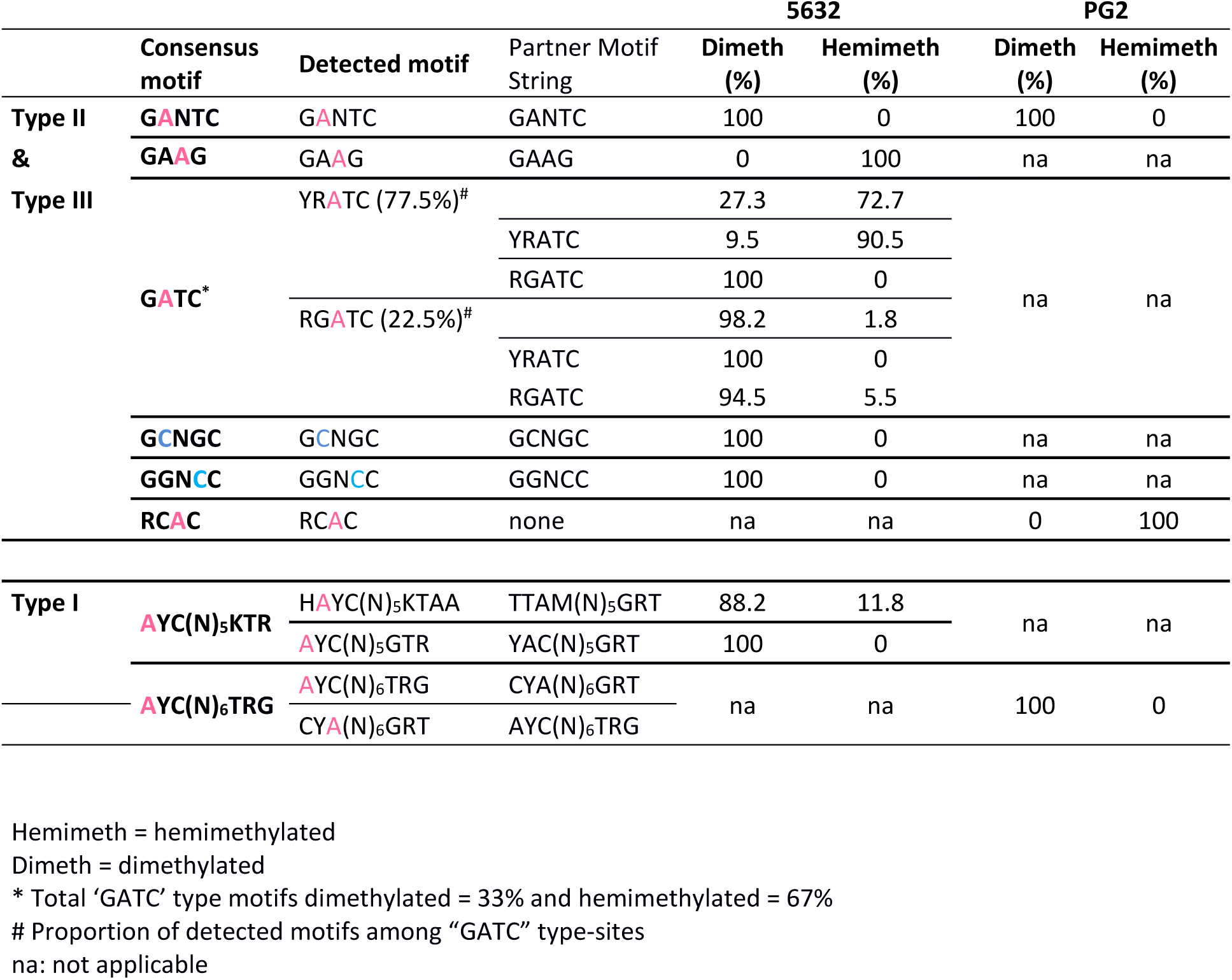
Methylation status of detected motifs in 5632 and PG2 *M. agalactiae* strains

Methylome data obtained with the panel of eight additional strains revealed the occurrence of both m6A and m5C methylated bases but not m4C, which remained specific to 5632 (Figure 1). The occurrence of m4C is not uncommon in prokaryotes, with m4C and m5C accounting for 20% and 5% of the modification types, respectively (27). Whether m4C may confer a particular property to 5632 is discussed later. Overall, this extended analysis identified a total of 19 methylated motifs, of which 11 had not been previously detected in 5632 and PG2: eight were assignable to Type I RM and eleven to Type II and III RM systems (Figure 1A). Of the detected motifs, 14 were associated with m6A modifications, with ten palindromic sequences that were methylated on both strands, while the other four imperfect palindromes (5’-GA^m6^TGC-3’, 5’-GA^m6^AG-3’, 5’-RCA^m6^C-3’, and 5’-GA^m6^GG-3’) were hemimethylated. Among those, two were Type III.

**Figure 1.**
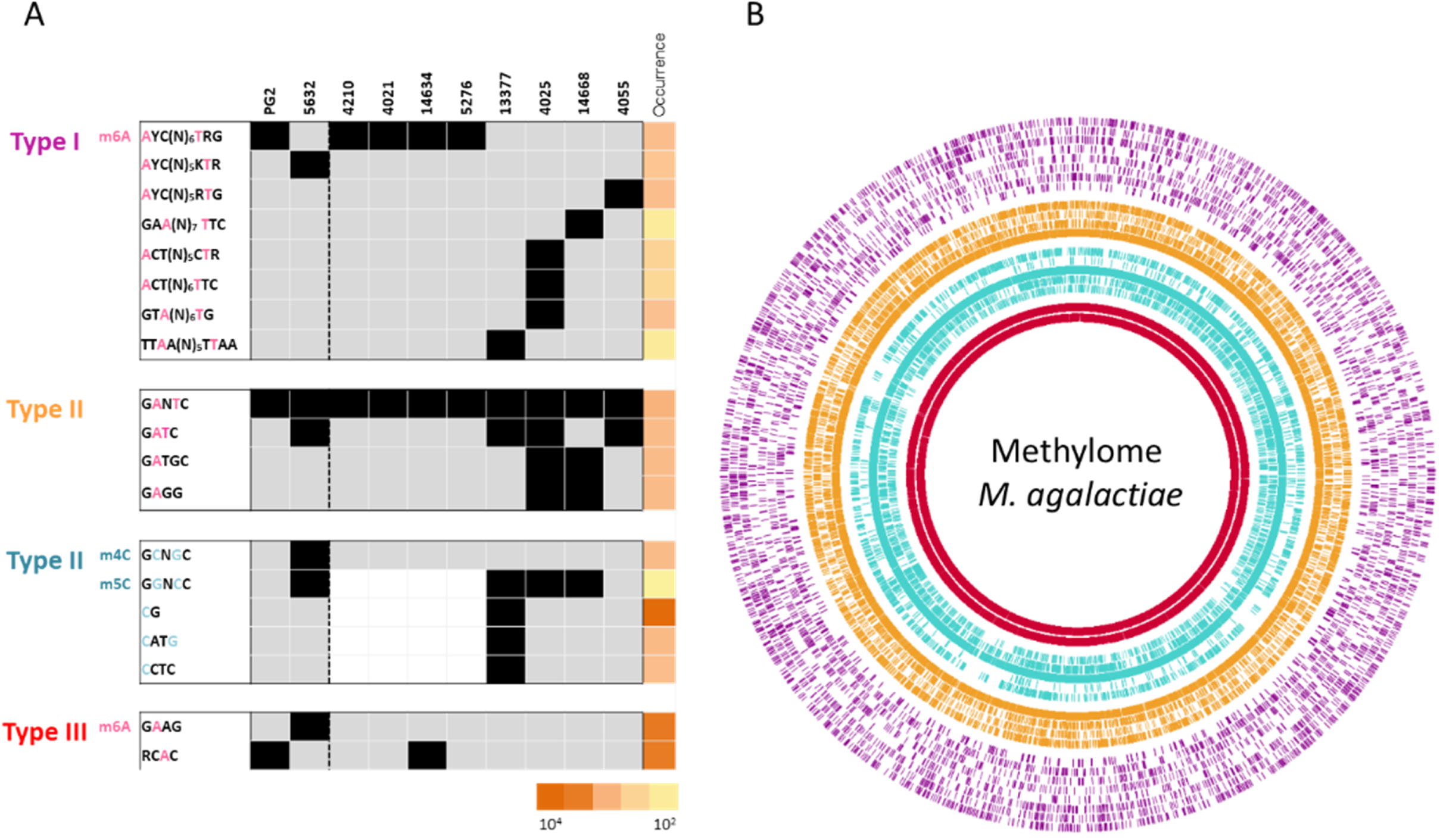
*M. agalactiae* epigenome. **(A)** Heatmap representing the occurrence and distribution of methylated motifs detected by SMRT and BS-sq analyses across 10 *M. agalactiae* strains. Motifs were grouped based on their assignment to Type I (purple), Type II (yellow for the Type II m6A methylated motifs and blue for the Type II m4C and m5C methylated motifs), and Type III (red) RM systems. The type of modification is also indicated: pink for adenosine methylation and blue for cytosine methylation. Strains not analysed by BS-seq are represented by a white box. The occurrence of methylated sites was evaluated *in silico* on the reference genome 5632 and is indicated by a column representing degrees of orange (the more intense the colour, the more the site is represented on the reference genome). **(B**) Circular plot representing the *in silico* position and occurrence of detected methylated motifs on the 5632 *M. agalactiae* reference genome. Methylated motifs were ordered as in the heatmap of panel A and grouped by Type and base modification. From outer to inner circle: Type I in purple, Type II m6a modification in orange, Type II m4C and m5C modification in blue, and Type III m6a modification in red.

Based on their methylome profiles, the panel of 10 *M. agalactiae* strains tested in this study could be divided into two main groups (Figure 1). The first had a methylome profile close to that of PG2, with a maximum of three distinct methylated motifs. Strains clustering in this group were further referred to as the PG2-like strains and included PG2, 4210, 4021, 14634, and 5276. They all shared the same Type I recognition motif (5’-A^m6^YC(N)_6_TRG-3’), in addition to the Type II 5’-GA^m6^NTC-3’ motif present in all *M. agalactiae* strains. PG2 and 14634 strains also possessed the 5’-RCA^m6^C-3’ Type III methylated motif. The second group of strains (5632-like strains) included 5632, 13377, 4025, 14668, and 4055 and displayed more complex methylome profiles, with up to eight different types of methylated motifs for strain 13377. One particular feature of this group was the presence of Type II m5C MTase genes, as revealed by BLAST analyses against the dcm REBASE database (24), with that of 4055 being truncated (Table S1). In agreement with this observation, m5C methylations were detected in all 5632-like strains except 4055 and PG2-like strains (Figure 1).

### *M. agalactiae* encodes sophisticated, variable, Type I RM systems

Type I RM systems are complex. In addition to restriction (R) and modification (M) subunits, Type I systems feature an additional specificity (S) subunit and work as multimeric complexes that recognise asymmetric sites. Typical Type I methylated motifs were detected in both 5632 and PG2 (Table 3) that consisted of a specific bipartite sequence separated by a 5N- and 6N-spacer, respectively. More specifically, in 5632, SMRT analysis identified four different, main motifs (Table 3) corresponding to six detected Type I methylated sequences (Table S3) and to the consensus sequence 5’–AYCN_5_KTR-3’ / 3’-TRGN_5_MAY-5’ with the target recognition domain (TDR) tandem TRD1 = AYC/GRT (Y = C or T and R = A or G) and TRD2 = KTR/YAM (K = G or T and M = C or A). Similarly, this analysis detected two different Type I motifs in PG2 corresponding to four methylated sequences and the consensus sequence 5’-AYCN_6_TRG-3’/3’-TRGN_6_AYC-5’, with TRD1 = AYC/GRT and TRD2 = TRG/CYA (Table 3 and Table S3). Type I specific methylated motifs were further detected in all *M. agalactiae* strains examined in this study (Table 3 and Figure 2). More specifically, strains of the PG2-like group, namely 4210, 4021, 14634, and 5276, all shared the Type I methylated motif 5’-AYC(N)_6_TRG-3’ found in PG2, while a greater diversity of sequences was observed for this motif in 5632-like strains. Notably, a single, specific Type I methylated motif occurred in all strains, except 4025, which had up to three different motifs (Figure 1).

**Figure 2.**
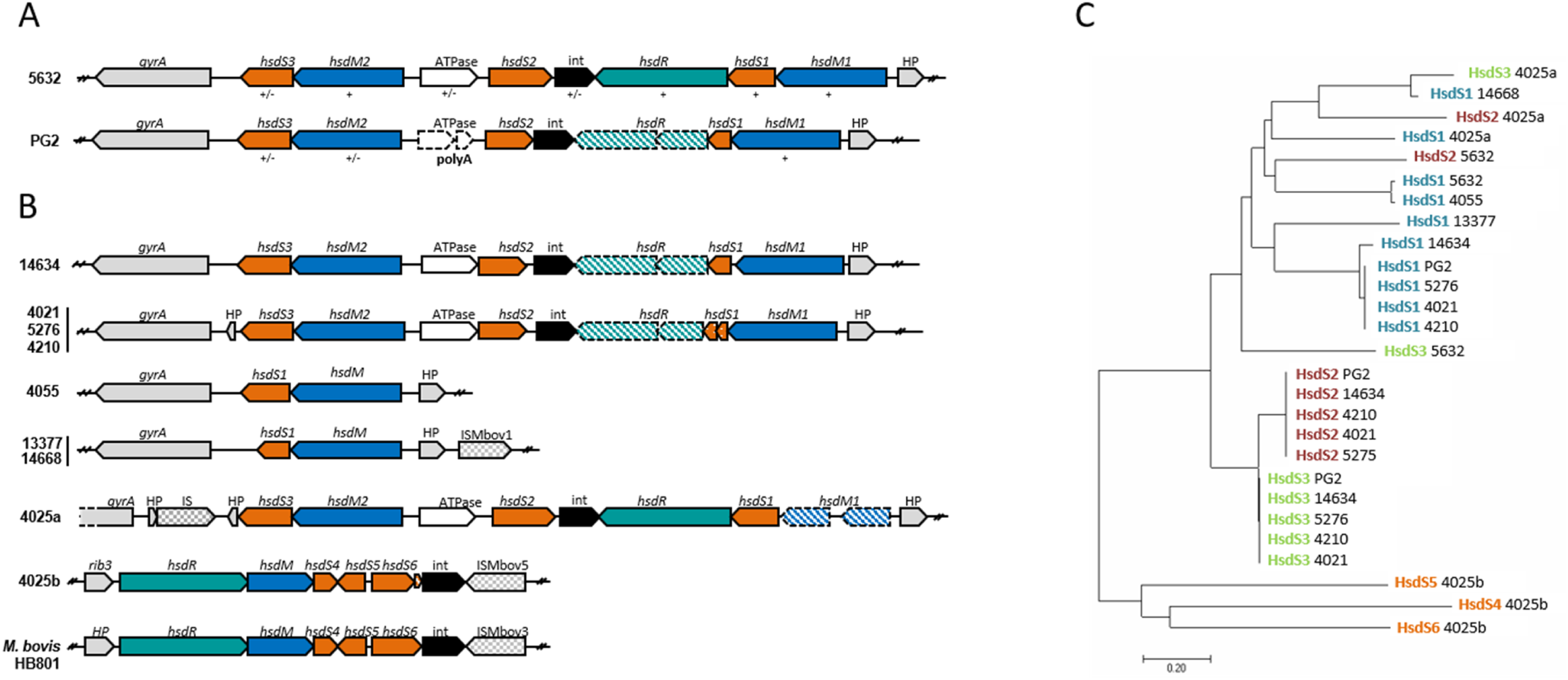
*M. agalactiae hsd* loci encode a Type I RM system. Schematic representing gene organisation of the *hsd* loci in **(A)** the two *M. agalactiae* reference strains 5632 and PG2 and **(B**) eight *M. agalactiae* field strains. Genes are indicated by an open arrow coloured based on the predicted function of their products (blue for methyltransferase, green for endonuclease, orange for DNA recognition subunit, black for integrase, and grey for other predicted gene). Pseudogenes are indicated by dotted lines and hatched genes correspond to truncated or size reduced genes. IS related transposase are indicated by grey squared arrows. **(C)** Evolutionary relationships of HsdS subunits. The evolutionary history was inferred using the neighbour-joining method based on a MUSCLE alignment and the optimal tree is shown (sum of branch length = 6.55524456). The tree is drawn to scale, with branch lengths in the same units as those of the evolutionary distances used to infer the phylogenetic tree. The evolutionary distances were computed using the Poisson correction method and are in the units of the number of amino acid substitutions per site. The analysis involved 27 amino acid sequences. All positions containing gaps and missing data were eliminated. There were a total of 36 positions in the final dataset. Evolutionary analyses were conducted in MEGA7 (67).

Several Type-I systems have been described in mycoplasmas at the gene level and were designated as host-specific determinant (hsd) systems based on the seminal work performed in *M. pulmonis* (36). In *M. agalactiae, hsd* loci were previously identified by BLAST analyses in strains 5632 and PG2 (18). As depicted in Figure 2A, their overall organisation and synteny are similar, and each contained genes encoding the restriction enzyme *hsdR* (*hsdR* is truncated in PG2, see below) and several copies of the DNA modification (*hsdM*) and specificity (*hsdS*) components. BLAST analyses showed that amino acid sequences of HsdM were highly conserved, while those corresponding to HsdS differed within and between the two strains (Figure 2A and Figure S2). This raised the possibility that HsdS variability might be responsible for the Type I DNA diversity observed in 5632 and PG2. Type I sequences may be recognised by different HsdS subunits whose abundance, turnover, or affinity for the sequence may vary. In agreement with this hypothesis, the relative proportions of the detected TRD diverged from their theoretical abundance estimated *in silico* (Table S3), ranging from 7 to 31% and 16 to 44% for 5632 and PG2 respectively (Table S3), while the proportion of different Type I detected motifs was comparable within each strain (63–69% in 5632 and 68–77 % in PG2; Table 3). In *M. pulmonis*, phase variation through TRD ‘shuffling’ by recombination of the different *hsdS* genes has been demonstrated (36). In this species, the Hsd system is composed of two distinct loci containing one *hsdR*, one *hsdM*, and two *hsdSs*, with up to 16 different profiles of *hsdS* genes that could be generated by DNA inversions (36). In *M. agalactiae*, the organisation of the *hsd* locus resembles that of *M. pulmonis* with two *hsdMs*, one *hsdR*, and three *hsdS* genes that occur as a single copy (Figure 2). Whether *M. agalactiae hsdS* genes also undergo recombination to generate *hsdS* variation is not known, but such an event may explain the variability of the Type I profile detected in both 5632 and PG2 strains. Interestingly, a potential recombinase (“int” in Figure 2) is encoded by the *M. agalactiae hsd* cluster, whose putative contribution toward DNA rearrangement within the locus remains to be addressed.

BLAST analyses further retrieved an *hsd* system in all *M. agalactiae* strains tested in this study (Figure 2B). While their overall composition and organisation were similar, variations across strains were observed. Most strains contained at least three *hsdS* copies, and two, *hsdS2* and *hsdS3*, were highly conserved among PG2-like strains (Figure 2 and Figure S2). These strains displayed the same Type I motif, but as it was degenerated, more than one base was permitted at a particular position of the target sequences (Figure 1A). This raised the question whether each *hds* subunit recognised a single sequence or whether there is a certain permissiveness of the subunits towards different sequences. A single *hsdS* gene copy was present in 13377 and 14668, whose respective Type-I motifs corresponded to a single, distinct Type I sequence (Figure 2). However, strain 4055, which also had a single *hsdS* copy, displayed a degenerate Type I sequence motif, but its mean score was among the lowest and the closest to the cut-off value, suggesting that this data may rather reflect a technical bias. At the other end of the spectrum, 4025 was equipped with six *hsdS* subunits distributed in two *hsd* loci: *hsd*4025a and *hsd*4025b (Figure 2B). Unsurprisingly, 4025 had the highest number of detected Type I methylation sites, with approximately 1900 sites for 1600 in 5632 and 1050 in PG2. SMRT analysis revealed that 4025 presented three Type I methylated motifs: TRD1 = ACT/RTC, TRD2 = ACT/CTT, and TRD3 = GTA/GT, with either five or six nonspecific nucleotides as spacers (Figure 1). Overall, these data suggested that strains equipped with a single HsdS/HsdM module tend to have a single, specific Type I methylated site and that the diversity of Type I methylated motifs correlates with the number of *hsdSs* within one strain.

To further address the impact of domain specificity on *M. agalactiae* methylation, SMRT-seq was performed with a 5632 variant, namely H1-2, in which the complete *hsd* locus was replaced by that of PG2 (13; see Materials and Methods). Methylome analysis of the 5632 H1-2 variant revealed the occurrence of the PG2 Type I methylation profile, with TRD1= AYC/GRT, TRD2 = TRG/CYA, and a 5N-spacer (see 5632 [hsd]_PG2_ in Table 3). Thus, replacement of the 5632 *hsd* locus by that of PG2 resulted in changing the Type I methylome profile of 5632 and supported that the *hsd* locus is responsible for Type I methylation, with variations of HsdS profiles being involved in modulating Type I targeted sequences.

The combination of multiple *hsdS* alleles with a potential recombinase gene in one locus provides a unique setting for *hsdS* sequence shuffling by DNA recombination, a phenomenon that, in turn, may drive recognition motif variations and dynamics in this species. In other mycoplasmas, Atack *et al*. (37) demonstrated that variation in the length of simple sequence repeats, such as AG[n], located in the HsdS coding sequence, was capable of mediating phase variation, thus leading to differential methyltransferase expression or specificity. Such repeats were not found in *M. agalactiae hsdS* genes, leaving gene ‘shuffling’ the best hypothesis for Type I variations.

*M. agalactiae hsd* clusters were always located at the same chromosomal position, downstream of *gyrA*, with the situation of 4025, which had an additional locus, *hsd*4025b, located elsewhere, downstream of the ribulose phosphate 3-epimerase *rib3* gene (Figure 2B). The organisation of this locus is similar to the *hsd* locus of *M. bovis* strain HB0801, with 93% identity, while the *hsd*4025a was closer to its counterpart in 5632. This suggested that *hsd*4025b was acquired from *M. bovis* by HGT, which was additionally supported by the occurrence of an *M. bovis* IS element, ISMbov5, at one end of the *hsd*4025b locus (Figure 2B).

Most *M. agalactiae* strains contained two *hsdM* genes, except for 13377, 14668, and 4055, which had only one (Figure 2B and Figure S2B). The extreme simplicity of the 13377 and 14668 *hsd* loci, which were only composed of an *hsdM*/*hsdS* module, the high similarity of their MTases with 5632HsdM1 (99.6% nt identity for 14668, 99.5% nt identity for 13377; Figure S2B), and the occurrence of an IS-related transposase (ISMbov1) near these loci suggested that these regions were subjected to evolutionary erosion (Figure 2B).

Finally, *hsdR* was disrupted in PG2, as well as in the 4021, 4210, 5276, and 14634 genomes (Figure 2B). This was due to the insertion of two nucleotides in a poly(A) region located in the middle of the gene that resulted in a premature stop codon (18 and this study) and a severely truncated product. This implies that *hsdR* is not required for Type I methylation to occur in *M. agalactiae*. Furthermore, a complex of HsdS and HsdM was shown to be sufficient in *E. coli* for the preferential methylation of the target sequence at adenine residues, while the three Hsd subunits, M, S, and R, were required for DNA cleavage (38). Truncation of *hsdR* also suggested that, in these strains, the Hsd system might be only capable of protecting DNA without restricting unprotected DNA. In mycoplasmas, polymeric tracts, such as poly(A) or poly(GA), are often prone to frequent insertion-deletion, a mechanism that could promote ON/OFF phase variation in the expression of the HsdR, as observed for the HsdS subunits in certain *Mycoplasma* spp. (37).

### Intra- and inter-strain diversity of the methylome produced by Type II RM systems in *M. agalactiae*

Type-II methylated motifs are usually short palindromes and were found in *M. agalactiae* with their occurrence, distribution, and sequences varying among strains, except for 5’-GA^m6^NTC-3’, which was detected in all strains. Based on REBASE prediction, this common motif can be associated with a Type II orphan MTase (Table 2) encoded by MAG6680 and MAGa7650 in PG2 and 5632, respectively, and by homologs in other strains (Figure 3). In *Caulobacter crescentus* and other bacteria (20, 39), the GANTC motif is also recognised by an orphan MTase that methylates the adenine. This MTase belongs to the cell cycle-regulated DNA MTase family (CcrM) and plays an essential role in regulating the cell cycle. Whether the MTase encoded by MAG6680 and its homologs has a similar function in *M. agalactiae* remains to be demonstrated.

**Figure 3.**
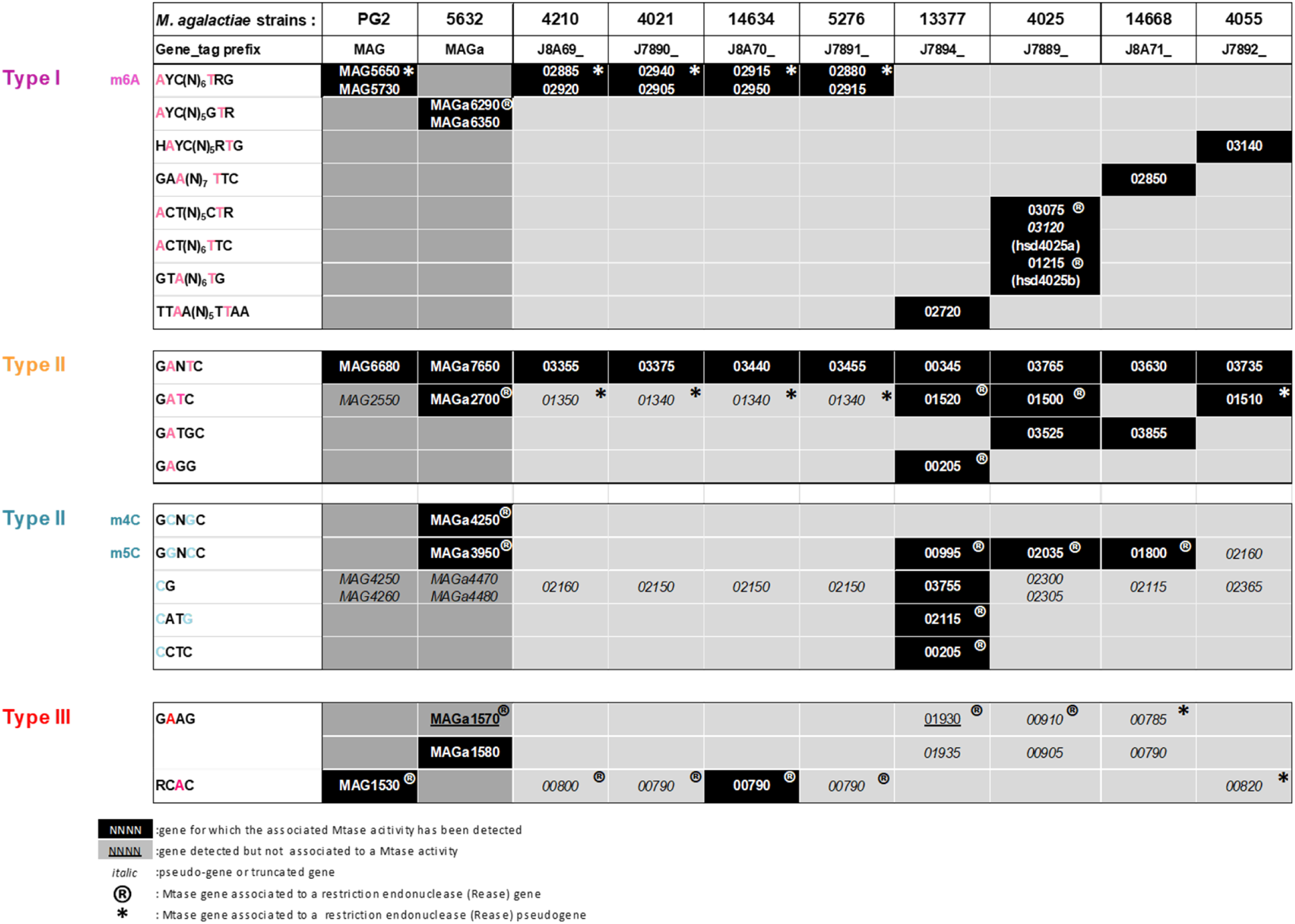
Methyltransferases assignation in *M. agalactiae* genomes. Methylated motifs detected by SMRT and bisulfite sequencing in ten strains of *M. agalactiae* and their associated methyl transferases. Motifs were grouped according to their assignment to Type I (purple), Type II (yellow for m6A and blue for m4C or m5C) and Type III systems (red). MTase gene name corresponds to the Genbank accession number generated after automatic annotation of the gene using the Prokaryotic Genome Annotation Pipeline (PGAP) from NCBI. The Gene_tag prefix is mentioned in the second line. Mtase genes that are associated with a cognate restriction endonuclease gene or pseudogene are indicated by a “R” or by an asterisk, respectively. Black boxes correspond to genes for which the associated Mtase activity has been detected. Underlined genes are those which have been detected genomes but for which no Mtase activity has been detected. Pseudogenes or truncated Mtase genes are shown in italics.

Two other degenerate Type II motifs, 5’-YRA^m6^TC-3’ and 5’-RGA^m6^TC-3’, were detected in 5632 but not in PG2. They were both assigned to a single MTase, MAGa2700, for which the predicted motif was 5’-GA^m6^TC-3’ (Table 3). Such an off-target methylation phenomenon has already been observed in *E. coli* for M.EcoKDam, a dam MTase involved in both chromosomal replication initiation and maintenance of genomic integrity (40). Close examination of the 5632 methylome further revealed that 77.45% of the GATC methylation corresponded to methylation of 5’-YRA^m6^TC-3’ (Y=C or T; R= A or G) and 22.55% to the 5’-RGA^m6^TC-3’ motif, with 72.7% of 5’-YRA^m6^TC-3’ being hemimethylated (Table 4). In all observed cases, this partial methylation occurred on one strand, usually the reverse strand, when the methylated site recognised was 5’-AA^m6^TC-3’ instead of 5’-GA^m6^TC-3’. The 5’-GA^m6^TC-3’ methylated motif was also detected in three other strains, namely 13777, 4055, and 4025, in which gene counterparts to MAGa2700 were identified (Figure 3). In the remaining strains, homologs to MAGa2700 were either detected as pseudogenes or totally missing, as in 14668 (Figure 3). In 5632, the MAGa2700 MTase is suspected to have a certain level of permissiveness regarding the recognition motif (see above), whereas in other strains the 5’-GA^m6^TC-3’ motif was the only detected sequence. Amino acid (aa) sequence alignment of products encoded by MAGa2700 and 13377, 4025, and 4055 homologs revealed two aa modifications out of 280 aa, in positions 22 and 97 of the active domain, which may explain the off-target methylation observed in 5632. Notably, 5’-GA^m6^NTC-3’ and 5’-GA^m6^TC-3’ were the most prevalent methylated motifs found among the Pacbio SMRT sequenced *Mycoplasma* sp. of the REBASE Genomes database. This supports that these epigenetic modifications play an important biological role in mycoplasmas.

A third Type II m6 methylated site, the 5’-GA^m6^TGC-3’ motif, was detected in two strains, 4025 and 14668, and BLAST against the REBASE database retrieved a cognate MTase gene only in the genome of these two strains (see Figure 3). These MTase coding sequences displayed 96% identity with a putative Type II N6-adenine DNA methyltransferase of *M. bovis* PG45 (MBOVPG45_0722), which recognised the 5’-GCA^m6^TC-3’ motif, the reverse complement of GATGC, and corresponded to the *Sfa*NI prototype of *Streptococcus faecalis* (REBASE database; 24). However, in 4025 and 14668, no cognate REase gene could be identified. Finally, the 5’-GA_m6_GG-3’ motif was only detected in the 13377 strain; the assignation of its corresponding MTase is discussed later.

The occurrence of Type II motifs with methylated cytosine, m4C or m5C, was also found in *M. agalactiae* but, as mentioned above, only in 5632-like strains. While all Type-II methylated motifs detected in 5632 were at least present in one other strain of this group, the 5’-GC^m4^NCG-3’ was an exception, and m4C modification was only found in this strain. Using REBASE, this motif was associated with the MAGa4250 MTase in 5632, which had no homolog in the other strains. Four other Type II motifs with methylated cytosine were detected, all with m5C. Of these, the 5’-GGNC^m5^C-3’ motif occurred at least in four strains, namely 5632, 13377, 4025, and 14668, but was not detected in 4055. This motif was associated with the MAGa3950 MTase of 5632, which homolog was encoded by 13377, 4025, and 14668 but was truncated in 4055 (Figure 3). The three other cytosine-methylated sites, 5’-C^m5^G-3’, 5’-C^m5^ATG-3, and 5’-C^m5^CTC-3’, were all detected in strain 13377 but not in other strains of our panel (Figure 1 and Figure 3). The case of the 5’-C^m5^G-3’ motif is addressed below, in the section dedicated to CpG methyltransferase. BLAST analyses using the REBASE database retrieved a cognate MTase gene for the other two motifs (Tables 3 and 5). Interestingly, BS-seq data showed that the 5’-C^m5^CTC-3’ was methylated on the cytosine, while its reverse complement, 5’-GA^m6^GG-3’, was shown by SMRT analysis to be methylated on the adenine and was further classified as a Type II motif (see above). The dcm MTases responsible for 5’-C^m5^CTC-3’ methylation encoded in 13377 by 13377_16.426 (Figure 3) had 92% identity with M.MboH1ORF688P of *M. bovis* Hubei-1 strain (prototype of the Type II, MnlI, CCTC(N)_7_). The MnlI prototype mentioned above can modify separate DNA strands on the same asymmetric target because of the two MTases present in the same operon: dcm M1.MnlI, which methylates the 5’-C^m5^CTC-3’ motif of the upper strand, and the dam M2.MnlI methyltransferase that targets the 5’-G^m6^AGG-3’ motif of the bottom strand (41). In *M. bovis*, M.MboH1ORF688P was initially annotated as a dam adenine-specific DNA methyltransferase and was recently reannotated as a Type II cytosine-5 DNA methyltransferase, probably recognising the CCTC motif. Interestingly, the non-mycoplasma MTase that had the most significant homology with MTase 13377_16.426 (63% identity) was M.Cal5972II of *Carnobacterium alterfunditum* strain pf4, for which the associated motif had been confirmed by Pacbio sequencing. This MTase was annotated as a m5 cytosine MTase, but Pacbio data suggested that it is responsible for the methylation of both the GA^m6^GG and T^m6^CCTCY motifs. The authors, in their comments, hypothesised that this second motif probably corresponds to the methylation of m5C in the complementary strand for which the modified base is not being called correctly (see REBASE pacBio data for M.Cal5972II). As for *C. alterfunditum*, our data strongly suggest that a single MTase, MTase 13377_16.426, is responsible for the methylation of the two asymmetric motifs 5’-C^m5^CTC-3’ and 5’-G^m6^AGG-3 (Figure 3). No dam MTase that could account for the m6A methylation of GAGG was detected in the 13377 genome, while the same motif was shown to be methylated differently. Such a situation is not unusual, and several studies have already shown the capacity of a given Mtase to change its target base specificity, such as *EcoR*V in *E. coli* or *Pvu*II in *Proteus vulgaris* (42, 43). For all Type II motifs with methylated cytosines described above, both cognate MTases and REases were identified (Figure 3), suggesting that all Type II motifs except the CG motif (see below) were associated with an RM system.

Finally, three Type II MTase genes were initially predicted *in silico* for which no corresponding methylated motif was identified, namely CDSH of 5632 and MAG3310 and MAG4030 of PG2 (Table 2). However, none of their corresponding products were detected by proteomic analysis, indicating that these genes were most likely not expressed under our laboratory conditions or at undetectable levels. While MAG3310 and MAG4030 are encoded by the chromosome, CDHS is a putative Type II MTase carried by an integrative conjugative element, ICEA, occurring in three copies in the 5632 genome but absent from the PG2 strain. Previous data suggested a complex interaction among identical ICEA copies integrated in the same genome, including downregulation of some components. The contribution of the putative CDSH MTase to the *M. agalactiae* methylome was then addressed in the PG2 strain, in which a single ICEA copy was introduced by HGT, namely the PG2-ICEA+ variant (Table S4). Comparison of SMRT data obtained with PG2-ICEA+ and WT PG2 showed no difference, confirming that the MTase CDSH was not active under laboratory conditions regardless of the number of ICEA copies encoded by one genome. However, mating experiment data suggested that CDSH was expressed in PG2 when introduced behind a constitutive promoter (see below).

### *M. agalactiae* Type III methylome profiles and their possible association with phase variable RM systems

Two different Type III methylated motifs were detected among the set of *M. agalactiae* genomes: 5’-GA^m6^AG-3’ and 5’-RCA^m6^C-3’. Both had a typical Type III motif, as they corresponded to dam methylation of short non-palindromic sequences. Although not initially predicted by REBASE, the 3’-RCA^m6^C-5’ motif was detected in PG2 (Table 2), in which a single Type III MTase gene, MAG1530, was predicted (Table 2). This motif was also detected in 14634, whose genome harboured an identical MAG1530 homolog, 14634_6.443 (Figure 3). No other strain was identified in our study as having a methylated RCAC motif, although four possessed MAG1530 homologs initially annotated as pseudogenes because of the occurrence of a frameshift mutation at the beginning of their coding sequences that generated a premature stop codon (Figure 4A; see below). Altogether, these data were in support of MAG1530 and 14634_1.453 as the best candidates to encode 3’-RCA^m6^C-5’ cognate MTases.

**Figure 4.**
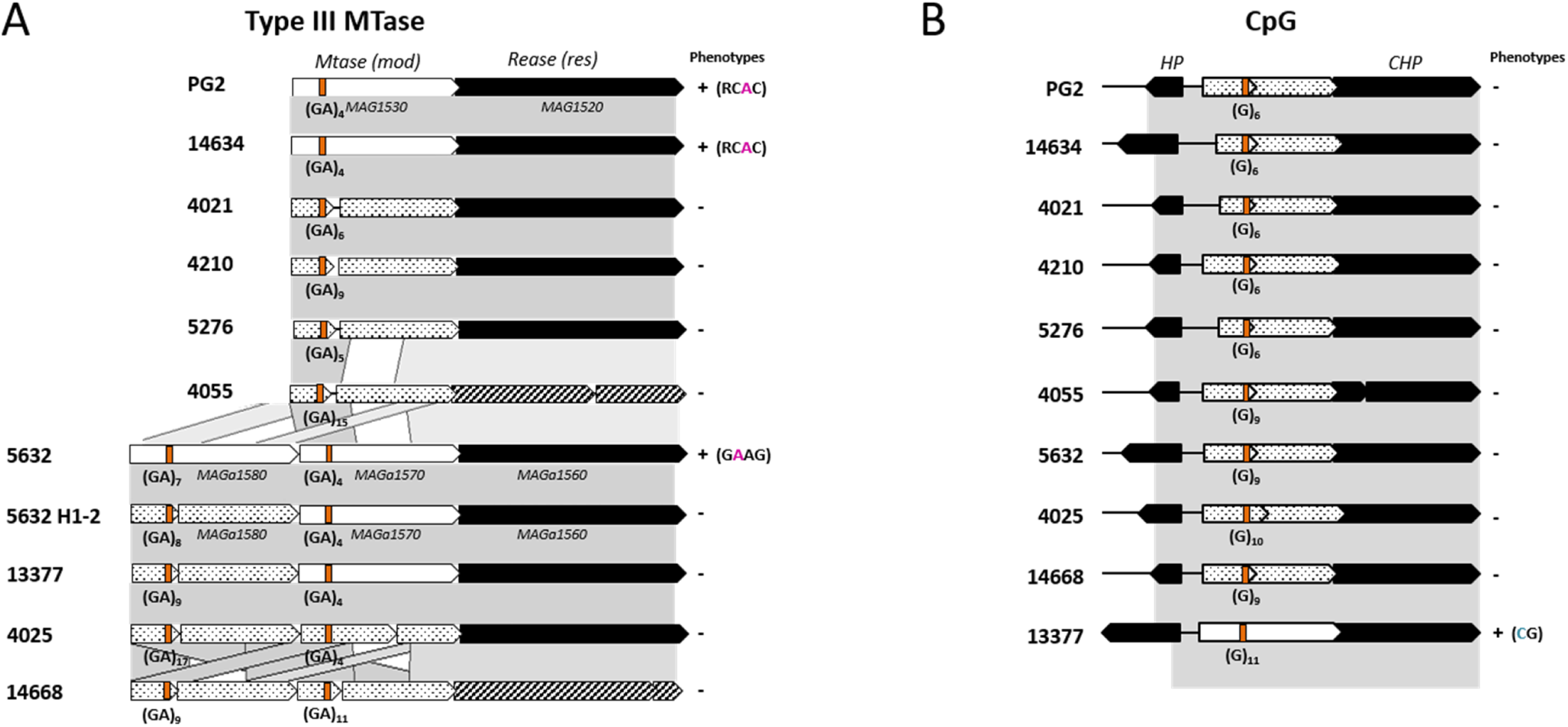
Type III RM loci and CpG methyltransferase in *M. agalactiae*. **(A)** Schematic representing the gene organisation and comparison between Type III RM systems identified in the ten sequenced *M. agalactiae* strains and in the 5632 H1-2 variant. Genes encoding mod methyltransferase (MTase) and restriction endonuclease res (REase) are indicated by white and black open arrows. Genes corresponding to truncated MTases are represented by open arrows filled with dots, and those corresponding to truncated REases are hatched. Poly(GA) tracts ([GA]) present in gene sequences are shown in orange, their length was validated by Sanger sequencing (see Materials and Methods) and is indicated below the orange box. Detection of the associated methylated motifs, the ‘phenotype,’ is specified on the right by a ‘+.’ **(B)** Schematic representing the CpG MTase gene across the ten *M. agalactiae* strains. CpG MTase genes are indicated by white arrows filled with dots when the gene is truncated. Black open arrows represent unrelated flanking genes. The poly(G) ([G]) tracts present in gene sequences are shown in orange, and their length was validated by Sanger sequencing (see Materials and Methods) and is indicated below the orange box. Detection of the associated CG methylated motifs, the ‘phenotype,’ is specified on the right by a ‘+.’

In the six strains considered above, MAG1530, 14634_1.453, and their homologs each occurred as a single copy (Figure 4A); however, these genes had strong similarities with two other CDSs of 5632 (64 to 70% identity), MAGa1570 and MAGa1580, which were also predicted as Type III MTases (Table 2). These were organised as a tandem in 5632 as in three other strains, 13377, 4025, and 14668, and shared 64.8% identity, suggesting that gene duplication was followed by genetic drift in these strains. Interestingly, no methylated GAAG motif could be detected in the 5632 H1-2 variant, in contrast to the 5632 parental strain (Table 3). Close examination of the 5632 H1-2 genome revealed a frameshift mutation located at the beginning of the MAGa1580 coding sequence that generated a premature stop codon (Figure 4A). Similar frameshift mutations were detected in 13377, 4025, and 14668, a finding that agreed with these strains having no methylated GAAG motif. Altogether, these data implicated MAGa1580 as the GAAG cognate MTase, but no methylated motif could be assigned to MAGa1570MTase. Notably, whether they occur as a single copy or as a tandem, these MTase genes were always detected upstream of the REase gene, with which they most likely formed an operon (Figure 4A), thus constituting a Type III RM system.

Type III RM systems are widespread in bacterial genomes and particularly in pathogens, in which they modulate genome methylation profiles by switching ‘on-off’ the expression of their MTases, an intra-clonal mechanism known as phase variation (44). Furthermore, Type-III N6-adenine DNA-methyltransferase encoding genes, known as *mod* genes, were previously described in mycoplasmas by Attack et al. (44) and more specifically in the *M. agalactiae* 5632 strain. In REBASE, these have been designated as M1 and M2.Mag5632ORF1570P and correspond to MAGa1570 and MAGa1580 of 5632, respectively, based on NCBI annotation. The features shared by *mod* genes include: (i) conserved 5’ and 3’ domains, (ii) conserved DPPY and FXXGXG Type III methyltransferase motifs required for function (44), and (iii) a variable central region encoding the TRD, which is responsible for the recognition of sequences methylated by the Mod protein. Upstream of this sequence, Atacks et al. (44) identified a tract of [GA] repeats in which the variation in number was responsible for phase variation events. Our study showed the occurrence of a similar tract of [GA] repeats in the repertoire of *M. agalactiae mod* genes, with the number of repeats varying within and among strains from 4 to 15 (Figure 4A).

In six strains, the *mob/res* locus was composed of a single MTase gene (MAG1530), followed by its cognate REase (MAG1520). In this group, Type III 3’-RCA^m6^C-5’ methylated sites were only detected in PG2 and 14634, an observation that correlated with the size of their poly(GA) being equal to 4 (Figure 1, Figure 4A). In the other strains, homologs to MAG1530 were truncated due to a change in the length of the *mod* [GA]_n_ repeated region that resulted in a premature stop codon. The *mob/res* locus was more complex for strains 5632, 13377, 4025, and 14668, in which it was composed of two MTases in tandem, namely MAGa1580 and MAGa1570 (5’ to 3’) for 5632, followed by one REase (MAGa1560; Figure 4A). In these strains, MAGa1580 and MAGa1570 homologs often occurred as pseudogenes due to frameshift mutations occurring in the *mod* [GA]_n_ region. Therefore, as shown in Figure 4A, all homologs to MAGa1580 were expected to produce truncated products due to the size of their [GA]_n_ that generated a premature stop codon. A similar situation occurred for MAGa1570 homologs of strains 4025 and 14668 and suggested that both MTase genes are subjected to phase variation in expression due to spontaneous insertion of [GA] in a poly(GA) region located at the beginning of the structural gene. The detection of the methylated motif 5’-GA^m6^AG-3’ only in strain 5632 and the absence of methylation of this very same motif in the 5632 H1-2 variant and other strains (Table 3) implicated MAGa1580 as the cognate MTase of the GAAG motif. The overall gene organisation of this cluster suggested that the two MTases were necessary for methylation to occur, knowing that Type III MTases are usually composed of a dimer of Mod subunits, where one Mod subunit recognises DNA, while the other Mod subunit methylates the target adenine (45).

Overall, differences in *mod* [GA] repetitions combined with the analysis of the methylome of all *M. agalactiae* strains and the 5632 H1-2 variant confirmed that the variations observed in this specific region had direct repercussions on the methylation status of mycoplasma genomes and reinforced the hypothesis that in *M. agalactiae*, Type III systems are regulated via phase variation, as observed in several other bacterial pathogens (46, 47).

### Detection of an active CpG methyltransferase in *M. agalactiae*

BS-seq data revealed the occurrence of about 20.10^3^ 5’-C^m5^G-3’ motifs in a single strain of the panel, namely in strain 13377. Such modification, termed CpG methylation, is common in eukaryotes, in which it plays an essential role in controlling gene expression. In contrast, prokaryotic CpG methyltransferases have been exclusively described in the class Mollicutes, which includes *Mycoplasma* spp. (48–50). In the AT-rich Mollicutes genome, the ‘CG’ motif is underrepresented, as observed in all *M. agalactiae* sequenced genomes, with a ‘CG’ abundance ranging from 0.46 and 0.5 (calculation based on the observed/expected ratio described by Goto et al. [51]; Table S5).

Data collected here showed that strain 13377 possessed an active CpG MTase for which a coding sequence, namely 13377_9.1088, was identified by BLAST analysis against REBASE dcm databases (Table S1). This coding sequence displayed 40% identity with the MSssI and MmpeI CpG MTases found in two other Mollicutes, *Spiroplasma* spp. strain MQ1 and *Mycoplasma penetrans*, respectively (48, 49). *Spiroplasma* MSssI was the first prokaryotic MTase described that specifically and exclusively methylates the 5’-CG-3’ sequence (48). In this Mollicute, as in *M. agalactiae*, these CpG MTases methylate both strands, while their eukaryote counterparts preferentially generate hemimethylated DNA (52). This is the first description of a ruminant pathogenic mycoplasma strain harbouring an active CpG MTase.

Comparative genome analysis revealed that all *M. agalactiae* genomes contained CpG MTase homologs (Figure 3). A common feature of these coding sequences is that they all displayed a poly(G) region having between 6 and 11 ‘G’ located in their middle. As shown in Figure 4B, the length of this poly(G) correlates with many of these gene products being truncated due to a premature stop codon. Only one strain, namely 13377, possessed an entire and active CpG MTase gene. Frameshift insertions-deletions occurring in CpG MTase genes have been previously described for other *Mycoplasma* spp. Moreover, in *M. pulmonis* or *M. crocodyli* (49), these were shown to modulate the length of a poly(GA) region also located in the middle of the CpG MTase sequence, as observed above for the poly(G). These poly(GA) or poly(G) stretches are located just upstream of the conserved and active domain of the dcm MTase (Figure 4B). These data reinforced the hypothesis that expression and functionality of the Mycoplasma CpG MTase active domain are controlled by frameshift mutations occurring in these regions and are thus phase variable.

In *M. hyorhinis*, CpG MTases and a CG overlapping GATC MTases (5’-CGAT^m6^CG-3’) were shown to translocate into the nucleus of human cells and efficiently modify their cellular epigenetic profiles. The presence of an active CpG MTase may play a significant role in the survival, pathogenesis, and host adaptation of *M. agalactiae*, which is also capable of invading host cells (53). Whether frequent, spontaneous mutations occurring in their coding sequences is a stochastic mechanism modulating MTase activity within clonal populations is not known, but the conservation of this phenomenon across *Mycoplasma* spp. indicates an important role.

### Abundance and distribution of methylated sites along the *M. agalactiae* genome

To gain insight into the putative role of DNA methylation in *M. agalactiae* biology, the predicted genomic distribution of methylated motifs was compared to that detected by SMRT- and BS-seq analyses using 5632 and PG2. Data showed that all *in silico* predicted motifs were indeed methylated under our laboratory conditions. Motifs having m6A modifications, 5’-GA^m6^AG-3’and the 5’-RCA^m6^C-3’ were the most abundant in terms of the number of methylated sites across all tested *M. agalactiae* strains (Figure 1A and B). They were found every 100 bp, on average (about 10^4^ predicted sites in the 5632 genome of 1 Mb). Remarkably, the 5’-GA^m6^NTC-3’ motif was the only one present in all investigated *M. agalactiae* strains, with 2.4 × 10^3^ sites per 1 Mb of the 5632 genome. As for cytosine methylation, the 5’-C^m5^G-3’ was found every 100 bp on average for a 1 Mb genome and was thus the most represented m5C motif. All m4C and m5C methylated sites were present in an occurrence of about 10^3^/1Mb, except for 5’-GGNC^m5^C-3’, which was less abundant (278 predicted motifs in the 1Mb 5632-genome; Figure 1A).

The distribution of predicted methylated motifs was further analysed using DistAMo (Figure S2; 54), which determines motif distribution among genomes using codon redundancy to evaluate their relative abundance. This analysis revealed a bias towards the *oriC* and *ter* regions in 5632 for the 5’-GA^m6^TC-3’ (z-score: 1,988) and the 5’-GGNC^m5^C-3’ (z-score: 2,004) methylation motifs, respectively, while no bias was observed for PG2 (Figure S3). DistAMo analyses also pinpointed the genes involved in cell cycle regulation as being overmethylated in both strains 5632 and PG2. The *ssb* gene, coding for a single-stranded DNA binding protein, displayed a high number of GANTC methylated motifs and the *hit* gene, coding for a HIT-like protein, in which Type III methylated motifs 5’-GAA^m6^G-3’ and 5’-RCA^m6^C-3’ were significantly overrepresented in 5632 and PG2, respectively (Table S6). The function of the hit gene is unknown, but proteins containing HIT domains form a superfamily of nucleotide hydrolases and transferases. This observation, together with the distribution bias of some motifs, supports the hypothesis that *M. agalactiae* DNA methylation plays a role in cell cycle regulation, as demonstrated for *Caulobacter crescentus* and other bacteria (20, 39).

The number of methylated motifs was also calculated in 1 kb windows along the PG2 and 5632 chromosomes (Figure S2). No particular region or gene was undermethylated except for loci encoding the three copies of ICEA in 5632. Indeed, ICEA-encoded regions displayed a mean of 12.99 methylated motifs per 1 kb (out of a total 82 kb represented by the ICEs on the 5632 genome) for a calculated mean of 15.96/kb for the 5632 genome (ranging from 3 to 39). Whether this difference has a biological impact is not known.

### Role of *M. agalactiae* RM systems in HGT

In prokaryotes, RM systems are known to provide a defence against foreign incoming DNA. Mycoplasmas are notoriously difficult to manipulate genetically, and this may in part be attributed to RM systems that prevent the entry of unprotected DNA. For instance, the 5632 strain can be efficiently transformed by specific mycoplasma vectors carrying the selective gentamicin or tetracycline resistance marker (pMT85, pminiO/T; 33). Replacing these with the puromycin resistance gene (*pac*) designed by Algire and Lartigue (55) was successful in generating transformants using PG2 but not 5632. A close examination of the *pac* gene sequence revealed the presence of an Hsd-5632 recognition motif (5’-A^m6^YC(N)_5_KTR-3’) at the beginning of the gene (position 18/924, GenBank accession no. GQ420675.1). Therefore, when introducing the unprotected site in the 5632 WT, it was cleaved by the HsdR REase, resulting in the loss of the vector carrying the puromycin resistance marker. Furthermore, we were successful in obtaining puromycin-resistant transformants with the H1-2 variant, which corresponded to 5632 having acquired the PG2 Hsd system (see above, data not show). Unlike its parental strain 5632, the H1-2 variant contained the mutated *hsd*R gene of PG2, which was predicted to produce a truncated, non-functional REase in this strain. These data highlighted the role of the Type I *hsd* system in the maintenance of foreign DNA in *M. agalactiae* once acquired, depending on the source.

The flow of genetic information between bacterial cells, a process known as HGT, is a key factor of microbial evolution. In prokaryotes, RM systems play an important role in this process by modulating the delicate balance between genome evolution and functional preservation. We recently demonstrated the occurrence of two types of HGT in *M. agalactiae*: one involved the self-dissemination of an integrative and conjugative element (9, 11); and the other, designated as MCT, was the exchange of large chromosomal fragments by recombination events using an unconventional distributive conjugative mechanism (10, 12, 13, 56). A recurrent observation was that MCT presented an apparent polarity, with strain 5632 always identified as the recipient cell regardless of the mating partners. This was true even when conjugation was bypassed by PEG-induced cell fusion, suggesting that the asymmetry of the DNA transfer might be independent of the conjugative mechanism itself, and relied on cytoplasmic factors that allowed, or not, the survival and/or the incorporation of foreign DNA.

To test this hypothesis and evaluate the impact of DNA methylation on DNA transfer polarity, mating experiments were conducted using 5632 and PG2 complemented by either one of the six Type II/III MTase genes identified above as being specific of 5632. These were MAGa1570, MAGa1580, MAGa2700, MAGa3950, MAGa4250, and CDSH (see Tables 2 & 3). Complemented MTase activities were controlled by restriction assays using DNA extracted from complemented PG2 strains and commercialised restriction enzymes, when available, that targeted the motif corresponding to the tested MTases (Sau96I for MAGa3950, Fnu4HI for MAGa4250, and DpnI/DpnII for MAGa2700; see Materials and Methods; Figures 4 and S3). Restriction profiles confirmed that MAGa3950, MAGa4250, and MAGa2700 of 5632, once introduced into PG2, protect DNA from being restricted. These data demonstrated that MTases were expressed and active in complemented PG2 clones and further confirmed that the detected methylated motifs were correctly assigned to the MTases in 5632. Notably, no commercialised REase matched the motif recognised by MAGa1580. Complemented PG2 clones were individually mated with WT 5632, and their mating efficiencies were compared to those obtained in parallel with a pair composed of the parental strains, WT PG2 and WT 5632, as controls. The results were expressed as a percentage (Figure 5), where 100% represented no difference and thus no effect on HGT of the MTase expressed by the complemented-PG2. As shown in Figure 5, a decrease in mating frequency was observed with all complemented PG2, regardless of the MTase genes used. The most important effects were observed when using PG2 complemented with MAGa3950, MAGa4250, Mag1570, and CDSH, with a decrease of at least 70% of the mating frequency compared to the control. Of these, the MAGa4250 MTase had the most significant impact, showing a 95% decrease. The expression and activity of the ICEA’s CDSH MTase were not detected during the methylome analysis described above (see above), regardless of whether CDSH was carried by 5632-

**Figure 5.**
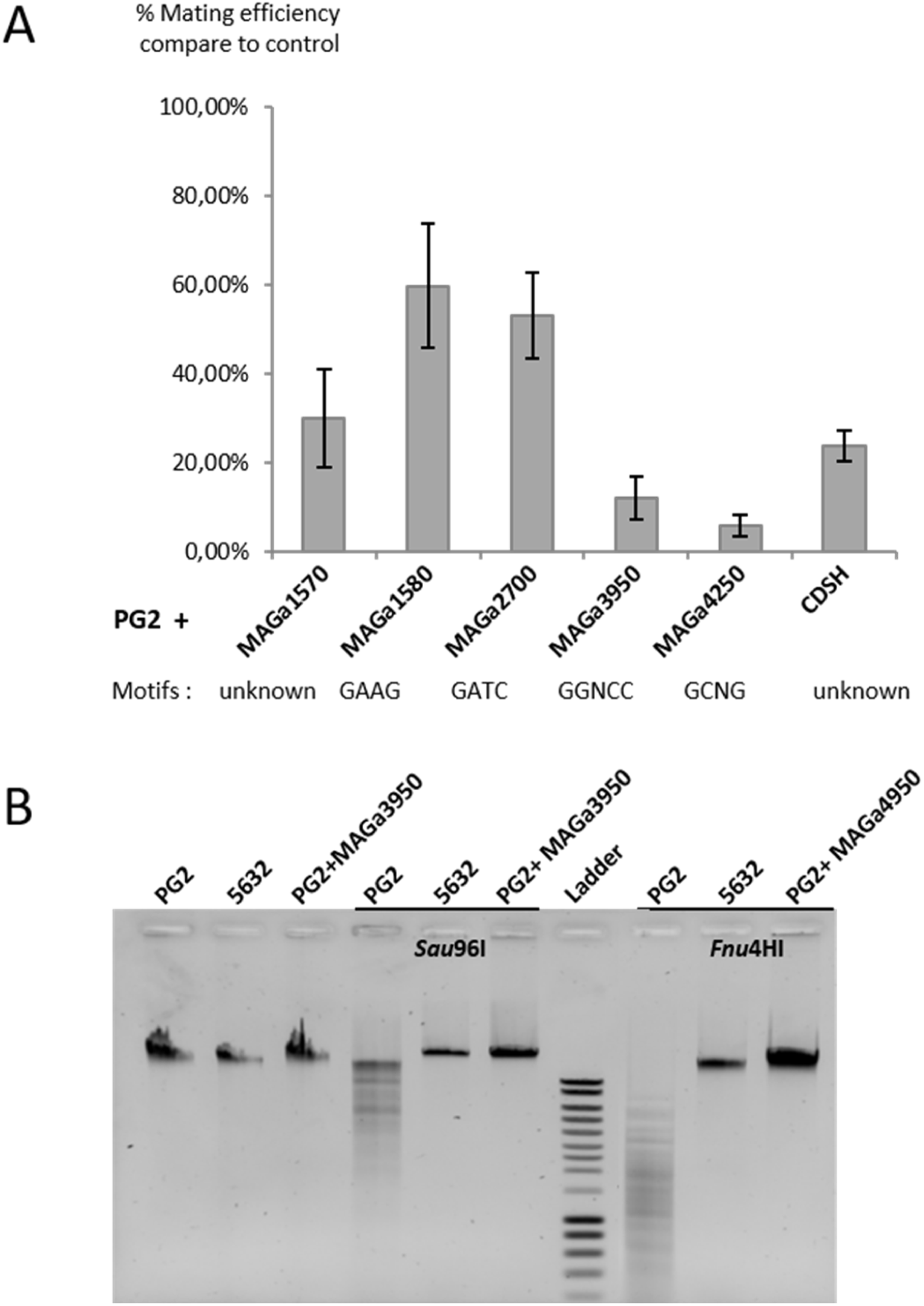
Contribution of *M. agalactiae* methyltransferases in mycoplasma chromosomal transfer. The PG2 strain was independently complemented by six MTase genes, absent in their active form in PG2 and originating from the 5632 strain, namely MAGa1570, MAGa1580, MAGa2700, MAGa3950, MAGa4250, and CDSH. The contribution of 5632 RM systems towards the acquisition of PG2 chromosomal DNA was addressed by comparing mating efficiencies obtained using 5632 and the individual complemented PG2 clones to that obtained with the non-complemented PG2 strain (see Materials and Methods). Mating frequency of the control corresponded to 6 ± 0.85 × 10^−8^ transconjugants/total CFU. For comparison purposes, results were expressed as a percentage of the ratio mating frequency assay/ mating frequency control, with values close to 100% representing no effect of the cloned MTase on DNA transfer on mycoplasma chromosomal transfer and those tending towards 0% having the greatest effect. Bar plots represent at least three biological replicates, with error bars as the standard deviation between replicates. When possible, complemented MTase activities were controlled by restriction assays as illustrated by agarose gel electrophoresis at the top of the bar plot. DNA extracted from 5632, PG2, and complemented PG2 strains were restricted by commercialised restriction enzymes *Sau96*I and *Fnu4H*I, which targeted the motif corresponding to the MAGa3950 and MAGa4250 MTases, respectively.

ICEA or PG2 ICEA+. However, when the same gene was introduced in PG2 behind a strong constitutive promoter (promP40), a reproducible, reducing effect towards the transfer of PG2 fragments into the 5632 genome was obtained. These data support our first hypothesis of a downregulation of CDSH when occurring in the context of ICEA and suggest that it may have a role in protecting DNA either from being transferred or incorporated into the recipient 5632 strain. Finally, the MAGa1580 and MAGa2700 gene products had a less significant impact, with only a 40% decrease in HGT efficacy.

Overall, these data experimentally support the general idea that one role of DNA methylation is to offer a protective barrier to HGTs. Thus, differences in RM system repertoires between *M. agalactiae* strains may explain the polarity repeatedly observed during chromosomal exchanges, with the 5632 strain always acting as the recipient strain. Furthermore, 5632 was the only strain harbouring the 5’-GCm4NGC-3’ motif and its cognate MTase MAGa4250, an epigenetic modification that is the most effective for protecting DNA (Figure 5). In our study, 5632 was shown to be the most methylated strain, with a total of 26,444 methylated sites (Table 3). It is also one of the richest in restriction endonucleases, with five associated REases (see Figure 3). These endonucleases may also play an important role in the mechanism of MCT by fragmenting the chromosomal DNA of the ‘donor’ strain, thereby facilitating its incorporation into the 5632 chromosome. Moreover, several studies have revealed that DNA restriction by REases generates products that appear to stimulate homologous and non-homologous recombination with the host genome (57) and thus may play a role in generating genomic diversity.

### Correlation between RM systems and the *M. agalactiae* mobilome

The interplay between bacterial RM systems and MGE is complex. Often considered as a barrier to invading MGE, RM systems can also stabilise MGE in cells by preventing infections by other competing MGEs (17). The association of MGEs and RM systems might result from an increased selection of RM systems facing foreign MGEs. As mentioned above, REases of these systems could also favour HGT by producing double-stranded DNA ends that are recombinogenic (17, 57, 58). In this case, genomes enduring more HGT and harbouring more MGEs would have more RM systems.

To evaluate the co-occurrence between MGEs and RM systems in mycoplasmas, the presence of MGEs in the 10 *M. agalactiae* genomes included in this study was investigated. For this purpose, for each strain, the repertoire of CDS related to mobile genetic elements (MGE), such as transposases commonly found in IS or integrase found in ICE and prophages, was analysed. MGE-associated CDSs were found in each genome, ranging from 18 to 113 per genome (Figure 6A, Table S7). Additionally, the minimal backbone of functional ICEs, with CDS5 and CDS17 encoding for TraE and TraG conjugation factors and CDS22 encoding for the ICE DDE-transposase (59), were retrieved in four strains (5632, 4025, 13377, and 14668; Figure 6A). Vestigial ICE forms were detected in nearly all strains, except 4055. Despite the paucity of prophage sequences in mycoplasmas, this study also retrieved a prophage in two strains (13377 and 14668). Thus far, *M. agalactiae* prophages have only been detected in strains isolated from ibex (60).

**Figure 6.**
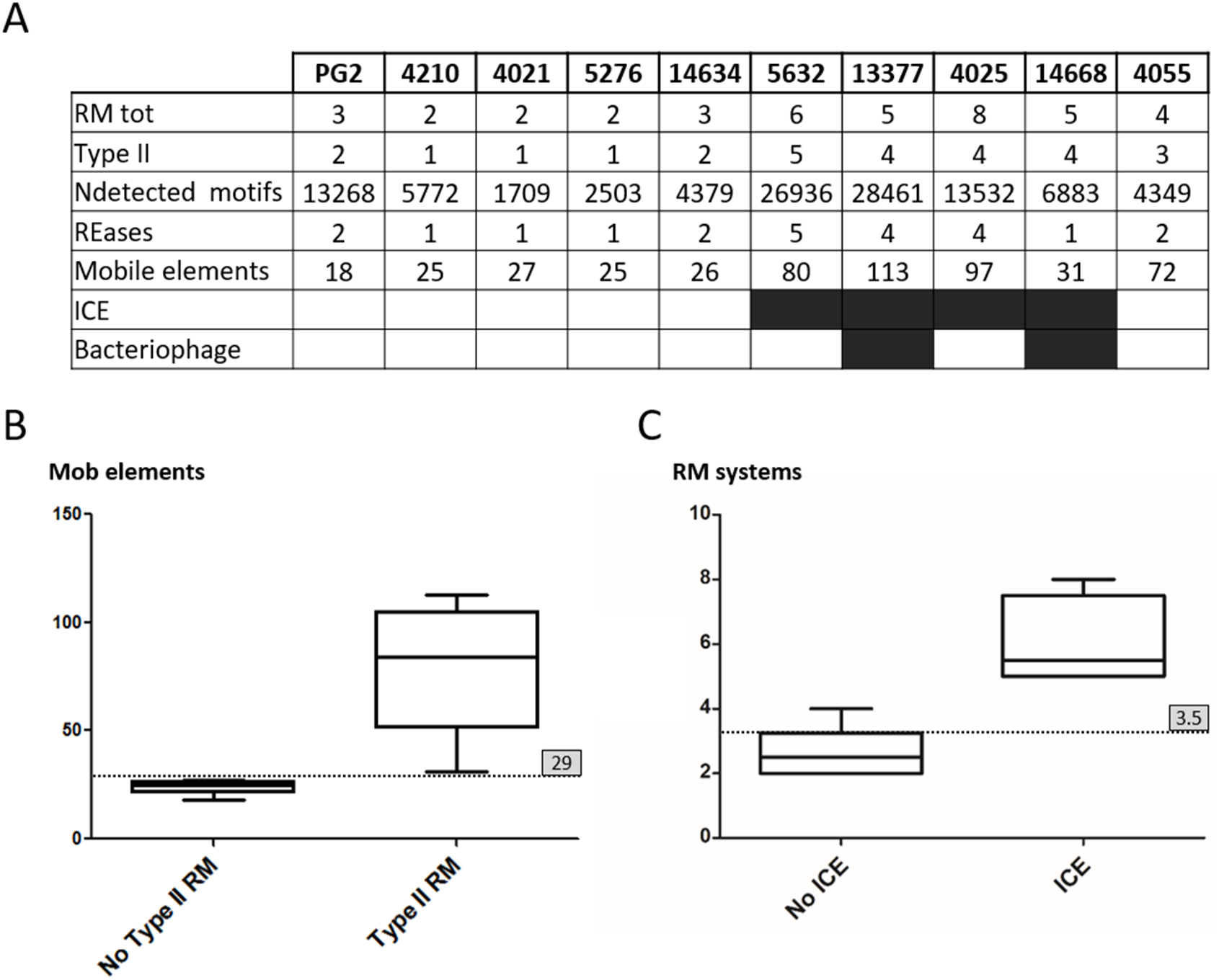
Correlation between horizontal gene transfer and restriction modifications systems in *M. agalactiae*. **(A)** Number of restriction modification systems (RMtot), Type II Rm systems (TypeII), methylated motifs (Ndtected motifs), restriction endonucleases (REase), and mobile elements detected in the ten strains of *M. agalatiae*. The number of mobile elements includes CDS annotated by RAST (30) as mobile elements, integrases, transposases and prophage elements and genes identified by blast against 5632 ICEA (e value superior to 10^−3^) as belonging to ICE elements. Based on previous work (59), elements were counted as ICE when at least the following three main components were present and complete: CDS5 and CDS17 encoding the conjugation factors TraE and TraG, respectively, and CDS22 encoding the integrase. **(B)** Boxplot showing the strong correlation between the presence of Type II active RM systems (excluding the 5’-GANTC-3’ methylation found in all strains) and the number of mobile elements (Pearson’s r = 0.78, p = 0.008) in *M. agalactiae* genomes. **(C)** Boxplot showing the very strong correlation between the presence of ICE elements and the number of RM systems in *M. agalactiae* genomes (Pearson’s r = 0.86, p = 0.001). For both boxplots, the dotted line represents the median; their values are indicated in the grey box on the right.

These data were used to evaluate the correlation between the presence of RM systems, the number of methylated sites detected (methylome), and the presence of MGE-related CDSs in the genomes of *M. agalactiae*. Our analysis revealed a strong correlation between active Type II RM systems and the presence of MGEs in *M. agalactiae* genomes (Pearson r = 0.78, p = 0.008). This correlation was also significant when all RM systems were considered (Pearson r = 0.71, p = 0.02; Figure 6B) and was even stronger when our analysis was restricted to ICE elements, which are an important contributor to HGT in mycoplasmas (9, 11; Pearson r = 0.86, p = 0.001; Figure 6C). Of note, the Type-II GANTC methylation that was broadly distributed across all tested strains and was suspected to be involved in cell cycle regulation, was not considered in this analysis.

Our results agreed with previous genomic studies that indicated a positive association between HGT capacity (based on MGE abundance in genomes) and the presence of RM systems, especially in larger bacterial genomes (17). This observation was made here with mycoplasmas, which are renowned for having some of the smallest bacterial genomes and is in opposition with the classical view in which RM systems contribute to bacterial immunity by protecting bacteria from exogenous DNA. To overcome this apparent contradiction, Rocha et al. (17) proposed three hypotheses.

The first was that orphan MTases encoded in MGEs promote genetic transfers by selfishly stabilising the presence of the element in the new host. Thus, genomes enduring more transfer would have more RM systems if they were carried by MGEs. This hypothesis is unlikely to explain the situation observed in *M. agalactiae*, since CDSH is the only MTase carried by ICEA in these bacteria; other RM systems detected in this study were all chromosomal. The second hypothesis formulated by Oliviera *et al*. (17) was that, within a clade, the abundance of RM systems would be linked to the selective pressure imposed by the abundance of MGEs. In this case, the selection pressure on RM systems should be higher for clades that endure frequent MGE infections. However, previous studies have shown that RM systems have limited efficacy and may not completely prevent the infection and spread of MGEs (61). This last scenario would result in a weak positive association between the transfer of genetic information and the abundance of RM systems and would contradict the data presented here (Figure 6), which are in favour of Rocha’s second hypothesis.

ICEs are intimately linked to HGT in mycoplasmas. They provide these bacteria with the capacity to conjugate, thus allowing them to exchange a large amount of chromosomal DNA via a conjugative, distributive mechanism designated as MCT. This process involves swapping large portions of the recipient mycoplasma genome by the donor counterparts via homologous and non-homologous recombination events (13). Thus, these exchanges of DNA materials may be facilitated by the presence of endonucleases, which would provide restriction breaks that stimulate recombination and promote HGT between cells. This scenario agrees with the last hypothesis of Rocha et al. (17), in which “R-M systems favor the transfer of genetic material between cells by generating restriction breaks that stimulate recombination between homologous sequences” and was consistent with the strong correlation observed between the abundance of REases and MGEs present in the *M. agalactiae* genomes (Pearson r =0.848, p = 0.0019; Figure 6B and C).

### Occurrence of *M. agalactiae* MTases outside the species

Whether MTases identified above were specific to *M. agalactiae* was addressed by BLASTP analyses against bacterial genomes available in current databases (Material and Methods). Furthermore, the five following MTase genes had no homolog outside of the Mollicutes class, which includes *Mycoplasma* spp., as well as *Ureaplasma, Spiroplasma*, and *Acholeplasma* spp.: the CpG methyltransferase, as previously described (see above), the Type III related MTases (MAGa1570, MAGa1580, and MAG1530), and the Type II MTase of 13377, 13377_964 (Table S8). Of the 11 Type II and III MTases identified in *M. agalactiae*, all were found in at least one other Mollicute, of a different species or genus and of a different phylogenetic group (Figure 7; Table S9). There was no correlation between their distribution and the phylogenetic cluster, the genome size, or the host of the species; however, this must be considered with care since the number of strains with sequenced genomes varies among Mollicutes. As shown in Figure 7, a remarkable observation is that homologs to these MTases were totally missing in eight *Mycoplasma* spp. For some, namely *M. cottewii, M. penetrans, M. pulmonis, M. arthritidis*, and *M. bovirhinis*, this may be due to the limited number of sequenced strains (n < 3). However, this explanation is not valid for *M. pneumonia* and *M. gallisepticum*, for which over 150 are available for each and unlikely for *M. genitalium* with 6 strains sequenced. Three *Mycoplasma* spp. that had different hosts all clustered in the Pneumoniae phylogenetic group. Remarkably, in these species no ICE was detected, and the *M. gallisepticum* genome carries an active CRISPR/Cas system (62), a bacterial adaptive immune system that protects from invasion by foreign DNA (63).

**Figure 7.**
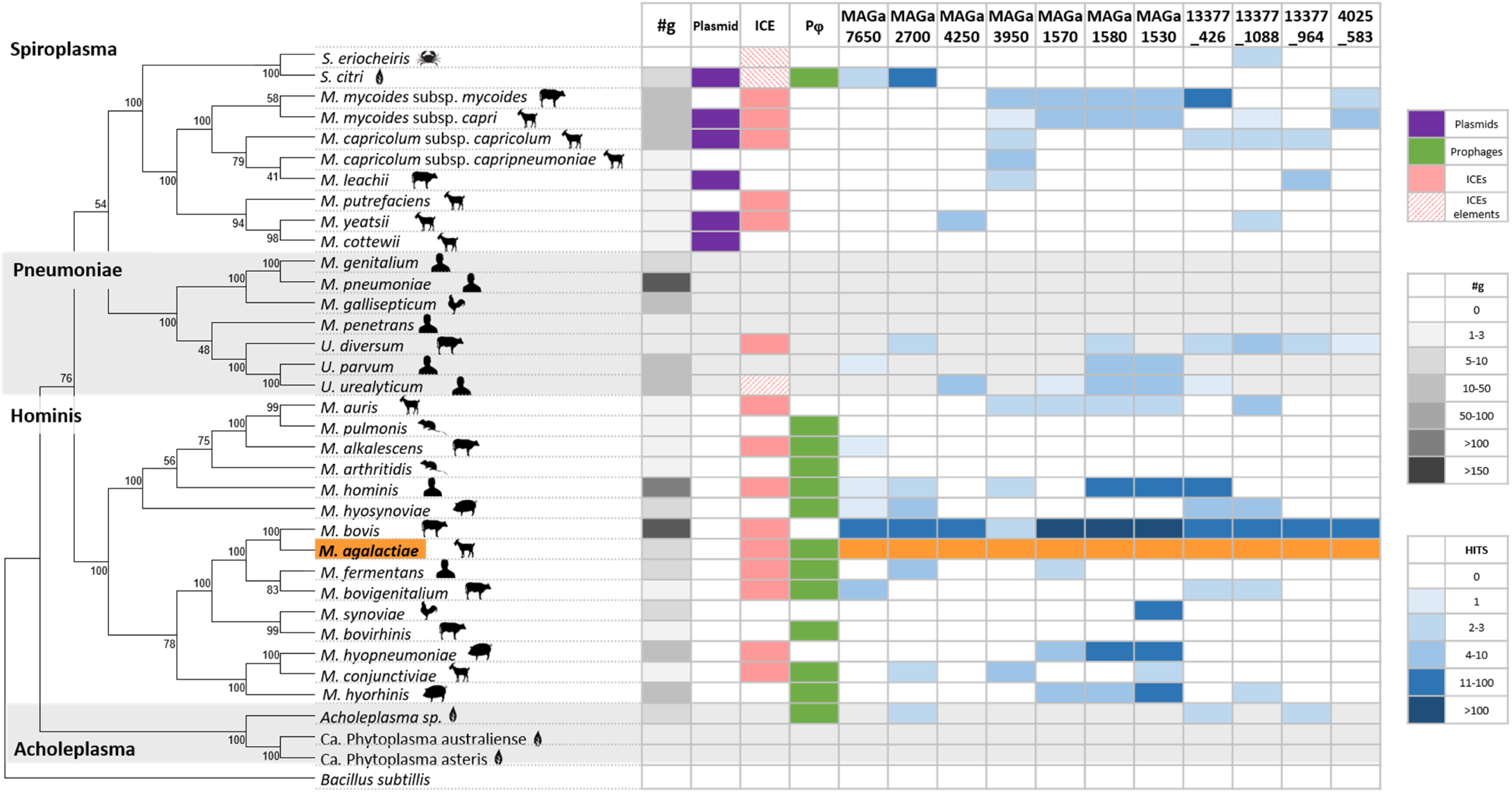
Distribution of active *M. agalactiae* methyltransferases among Mollicutes. Phylogenetic tree based on 16S ribosomal DNA sequences of 36 major representatives of Mollicutes aligned with clustalW (MEGA7; 67). The evolutionary history was inferred using the neighbour-joining method. The evolutionary distances were computed using the Poisson correction method. There were a total of 36 positions in the final dataset with a bootstrap of 500 replicates (percentage indicated next to each branch). *Bacillus subtilis* was used as an extra group. The main four phylogenetic groups are shown in the left. Our study model *M. agalactiae* is highlighted in orange. For each mollicute species, their host is symbolised next to their name. The proportion of sequenced genomes available in current databases are indicated in the ‘#g’ column by a grey gradient (the higher the number of genomes, the darker the grey). The presence in each species of plasmids, prophages, ICEs, or ICEs elements is represented by purple, green, pink, and hatched pink boxes, respectively. The occurrence and number of active *M. agalactiae* methyltransferases present in other mollicutes are indicated by blue boxes. They were determined by BLASTP analyses (NCBI Taxonomy results) on *M. agalactiae* methyltransferase protein sequences against mollicutes genomes available in current databases. The blue gradient represents the number of positive hits found after BLASTP analysis for each species (the higher the number of results, the darker the blue).

Finally, six MTases displayed significant homology outside of the Mollicutes, with some having up to 70% identity (coverage 100%) with CDS annotated as DNA methyltransferases in bacteria that are phylogenetically distant (Table S8). Altogether, our data indicate that *M. agalactiae* MTases identified in this study, and most likely their cognate REases when belonging to RM systems, frequently occurred outside of the species within the Mollicutes and outside of this particular class, most likely as a result of HGT.

## CONCLUSIONS

This study provides a comprehensive insight into the genome-wide methylome of several *M. agalactiae* strains at single-base and strand resolution using a combination of PacBio SMRT- and BS-seq. When combined with whole-genome analysis, this approach identified 19 methylated motifs associated with three orphan MTases and eight RM systems.

A single motif, the Type II G^m6^ANTC motif, was detected across all *M. agalactiae* strains tested here. This motif was associated with an orphan MTase having some similarities with the cell cycle-regulated DNA MTase family (CcrM) and is likely to play a role in basic, biological mycoplasma function(s). Two other Type II motifs were associated with orphan MTases, namely the G^m6^ATGC, which was detected in only two strains and for which there was no hint regarding a potential function, and the C^m5^pG which was only detected in one strain. In prokaryotes, CpG methylated motifs are exclusively found in the class Mollicutes, which includes *Mycoplasma* spp. The role of this methylation in mycoplasmas is not known, but studies conducted with *M. hyorhinis*, a facultative, intracellular pathogen of swine, suggested that this bacterium is able to deliver CpG methyltransferase into eukaryotic cells and selectively methylate the host genome, thus impacting its epigenome (50). Whether *M. agalactiae*, which is also described as facultatively intracellular, modulates its host epigenome remains to be elucidated. In *Mycoplasma* spp., CpG MTase is likely to be regulated at the population level, during clonal propagation, via high-frequency phase variation. Furthermore, a CpG MTase gene was found across all *M. agalactiae* strains that displayed a hot spot for spontaneous mutations typical of such a system. Orphan MTases were likely derived from RM systems by loss of the cognate REases at an early evolutionary stage history of the species (64). Some, such as the CcrM-like or CpG MTases, may be domesticated by the host chromosome to serve essential functions, thus becoming part of the core-genome of the species; others might have been further disseminated via HGT (16) as part of the variable pan-genome.

Most methylated motifs identified in this study were associated with the Type I *hsd* system described in other *Mycoplasma* spp., notably in *M. pulmonis*. In this species, the specificity of this system for a particular target sequence was shown to undergo phase variation following DNA shuffling within the *hsd* locus. The *hsd* genetic organisation and the diversity of the Type I m6A sequences identified in this study suggest that the same phenomenon is occurring in *M. agalactiae*, in addition to gene erosion and gene transfer. MTase expression of two other Type II and III RM systems identified here are also suspected to undergo phase variation based on frameshift mutations occurring in a homopolymeric tract, poly(G) or poly(GA). Such a genetic mechanism has been extensively described in mycoplasmas, in which it controls ‘on-off’ switching in the expression of a variety of products (65).

Based on their methylome profiles, the panel of *M. agalactiae* strains could be equally subdivided into two groups. One displayed only m6A methylated motifs together with a limited set of MTases and RM systems. The other group was more complex and diverse, including both methylated cytosine (m4C and m5C) and adenine (m6A) motifs, along with several RM systems and a CpG methyltransferase. The m4C is not a rare DNA modification in bacteria, but to our knowledge, this methylation has never been reported in mycoplasmas (26, 27). Here, m4C was only detected in the GCNGC motif of strain 5632, and in agreement with this finding, this strain was the only one equipped with the cognate MTase gene, namely MAGa4250. *M. agalactiae* strain 5632 can uptake and integrate chromosomal DNA from other strains via a conjugative, distributive mechanism designated MTC, which remains to be fully understood. This phenomenon was rarely observed with other strains, and when it occurred, it was at low frequency and in strains that were genetically similar to 5632. One hypothesis was that 5632 had the capacity to incorporate the incoming chromosomal DNA only once digested. In support of this hypothesis, our data indicated that cloning MAGa4250 into PG2 almost abolished the incorporation of PG2 chromosomal DNA into 5632, most likely by protecting the PG2 chromosomal DNA from being digested by the MAGa4250 Matse cognate REase. A search across the mycoplasma database (data not shown) retrieved a MAGa4250 homolog only in a few strains of *M. bovis*, a close relative of *M. agalactiae*, and two other ruminant *Mycoplasma* spp., *M. californicum* and *M. yeatsii*. Whether the rare occurrence of this MTase across Mollicutes is due to the low number of sequenced genomes for some species is not known. Thus far, the MAGa4250-RM system has been found in two distant phylogenetic groups in *Mycoplasma* spp. having the ruminant host as a common factor. This observation suggested that this system might disseminate via MTC across species sharing the same ecosystem, in addition to being a key player in this process. In *M. agalactiae*, RM systems may play a central role on MCT and on the polarity of DNA transfers in offering a delicate balance between the protective effect of MTases and the facilitation of DNA incorporation into the host chromosome by REases.

Overall, several superimposed genetic events may participate in generating a dynamic epigenome landscape within and among *M. agalactiae* strains. These occur at loci encoding orphan MTases or RM systems and include (i) DNA shuffling and frameshift mutations that affect the MTase and REases content of a clonal population during propagation and (ii) gene duplication, erosion, and transfer that modulate MTase and RM repertoires across the species. In turn, the versatility of these systems may contribute to regulating essential biological functions at both cell and population levels, and they may be key in mycoplasma genome evolution and host-adaptability by controlling gene flow among cells.

## SUPPLEMENTARY DATA

**Table S1**. BLAST analysis against dcm REBASE database

**Table S2**. Primers used for PCR and Sanger sequencing

**Table S3**. Percentage of methylated TRD couples corresponding to Type I RM systems detected by SMRT sequencing compared to *in silico* motif abundance in 5632 and PG2 strains

**Table S4**. Methylated motif detected in PG2 and PG2 ICEA+ variants

**Table S5**. CG abundance *of M. agalactiae* tested strains

**Table S6**. Over- and under methylated genes based on DistAMo analysis in *M. agalactiae* 5632 and PG2 stains

**Table S7**. Mobilome of the ten sequenced *M. agalactiae* strains

**Table S8**. BLASTP results for *M. agalactiae* active methyltransferases against bacteria other than the Mollicutes class

**Table S9**. BLASTP hits obtained for *M. agalactiae* active methyltransferase against the current Mollicutes genome database.

**Figure S1**. Patterns obtained after agarose gel electrophoresis of restricted DNA extracted from PG2 strain complemented with methyltransferases originating from the 5632 strain

**Figure S2**. Evolutionary relationship of HsdS, HsdM, and HsdR of the ten *M. agalactiae* tested strains

**Figure S3**. Distribution of *M. agalactiae* methylated motifs along the 5632 and PG2 chromosomes

## ACCESSION NUMBERS

Raw data and assembled genomes generated for this study have been submitted to NCBI (Bioproject PRJNA717738): SRR14651725, SRR14651724, SRR14651721, SRR14651727, SRR14651726, SRR14651723, SRR14651722, SRR14651720, SRR14651728, SRR14651719, SRR14651675, SRR14651674 for SMRT-seq raw data and SRR14329691, SRR14329692, SRR14329205, SRR14329203, SRR14329202,SRR14329204 for BS-seq raw data in the SRA database. Assembled and annotated genomes have been submitted to the GenBank database under the following accession numbers: JAGJTT000000000, JAGJTS000000000, JAGJTR000000000, JAGJTQ000000000, JAGJTP000000000, JAGRRE000000000, JAGJTO000000000, and JAGJTN000000000.

## ACKNOWLEDGMENTS

This work was performed in collaboration with the GeT core facility, Toulouse, France (http://get.genotoul.fr), supported by France Génomique National infrastructure, and funded as part of the “Investissement d’avenir” program managed by Agence Nationale pour la Recherche (contract ANR-10-INBS-09) and by the GET-PACBIO program (” Programme operationnel FEDER-FSE MIDI-PYRENEES ET GARONNE 2014-2020”). We are grateful to the Genotoul bioinformatics platform Toulouse Midi-Pyrenees, France, and the Sigenae group for providing support and storage resources. This manuscript was proof-read by a professional service.

