## Supplementary material for "Impacts of *Mycoplasma agalactiae* restriction-modification systems on pan-epigenome dynamics and genome plasticity": SupDataLegends.pdf

**Table S1**. BLAST analysis against dcm REBASE database

**Table S2.** Primers used for PCR and Sanger sequencing

**Table S3.** Percentage of methylated TRD couples corresponding to Type I RM systems detected by SMRT sequencing compare to *in silico* motif abundance in 5632 and PG2 strains

**Table S4.** Methylated motif detected in PG2 and PG2 ICEA+ variants

**Table S5.** CG abundance *of M. agalactiae* tested strains

**Table S6.** Over- and under-methylated genes detected by DISTAMO in *M. agalactiae* 5632 and PG2 strains

**Table S7.** Mobilome of the ten sequenced *M. agalactiae* strains

**Table S8.** BLASTP results for *M. agalactiae* active methyltransferases against bacteria other than the Mollicutes class

**Table S9.** BLASTP hits obtain for *M. agalactiae* active methyltransferase against current Mollicutes genomes database.

**Figure S1. Agarose gel electrophoresis of DNA restrictions of PG2 strain complemented with methyltransferases originating from the 5632 strain.**

Complemented activities were controlled by restriction assays when commercial restriction enzymes were available as illustrated in the agarose gel electrophoresis presented here. The DNA extracted from 5632, PG2 WT and complemented PG2 strains were restricted by commercialized restriction enzymes *Sau96*I , *Fnu4H*I, *Dpn*I and *Dpn*II which targeted the motif corresponding to the MAGa3950 and MAGa4250 Mtases and MAGa2700 (for both *Dpn*I and *Dpn*II), respectively**.** *Dpn*I and *Dpn*II recognize the same sequence but have different methylation sensitivities. DpnI will only cleave fully-adenomethylated dam sites and hemi-adenomethylated dam sites 60X more slowly. *Dpn*II cleave dam sites that lack adenomethylation and is blocked by complete dam methylation.

**Figure S2. Evolutionary relationships of HsdS, HsdM, and HsdR of the ten *M. agalactiae* tested strains**

Neighbor-Joining phylogenetic trees of the several *M. agalactiae* Hsd subunits. Proteins were aligned with MUSCLE implemented in MEGA 7. The phylogenetic tree was generated using the software MEGA7 (72). The tree is drawn to scale, with branch lengths in the same units as those of the evolutionary distances used to infer the phylogenetic tree. The evolutionary distances were computed using the Poisson correction method and are in the units of the number of amino acid substitutions per site. All positions containing gaps and missing data were eliminated. **(A)** Evolution history of all HsdS subunits detected in the ten *M. agalactiae* tested strains (strain name and sububunit number as assigned in Figure 2). The optimal tree with the sum of branch length = 6.55524456 is shown. The analysis involved 27 amino acid sequences and there were a total of 36 positions in the final dataset. **(B)** Evolution history of all HsdM subunits detected in the ten *M. agalactiae* tested strains (strain name and sububunit number as assigned in Figure 2). The optimal tree with the sum of branch length = 1.46693434 is shown. The analysis involved 18 amino acid sequences and there were a total of 418 positions in the final dataset. **(C)** Evolution history of all HsdR subunits detected in the ten *M. agalactiae* tested strains. The optimal tree with the sum of branch length = 1.41761699 is shown. The analysis involved 8 amino acid sequences and there were a total of 886 positions in the final dataset.

**Figure S3. Distribution of methylated motif along *M. agalactiae* 5632 and PG2 chromosomes**

**(A)** Circular plot representing the distribution of methylated motifs calculated per 1kb windows along the PG2 and 5632 chromosomes. From outer to inner circle for 5632 genome (grey circle) : GAAG (green), GANTC (dark blue), GATC (red), Type I methylated motifs (orange), GCNGC (purple), GGNCC (light blue). Integrative and conjugative elements (ICE) are represented by purple box in the grey outer circle representing the 5632 chromosome. From outer to inner circle for PG2 genome RCAC (green), GANTC (blue) and Type I methylated motifs (orange). (B) All methylated motifs distribution analyzed by Distamo (59) for 5632 and PG2 *M.agalactiae* chromosomes. The oriTer bias z-score corresponds to the bias of motif abundance on the oriC proximal half or the chromosome compared to the ter-proximal half of the chromosome. A positive z-score above 2 indicates a significantly higher abundance of the motif on the oriC-proximal half. A z-score below -2 indicates a significantly higher abundance on the ter-proximal half. The replichore bias z-score corresponds to the motif abundance on the right replichore compared to the left replichore. A positive z-score above 2 indicates a significantly higher abundance of the motif on the right replichore. A z-score below -2 indicates a significantly higher abundance on the left replichore.
