## Supplementary material for "Impacts of *Mycoplasma agalactiae* restriction-modification systems on pan-epigenome dynamics and genome plasticity": TableS1.pdf

**Table S1 : BLAST analysis against dcm REBASE database**

| Strain | Mnemonic | Homologs | | Predicted motif | Size (nt) | Identity (aa) |
|---|---|---|---|---|---|---|
| **4210** | 2.388 | M.MagPG2 | MAG3310P | uknown | 804 | 99% |
| | *2.486* | *M.Mag5632* | *MAGa4470* | *CG* | *426* | *96%* |
| | *2.487* | *M.Mag5632* | *MAGa4480* | *CG* | *747* | |
| **4021** | 1.360 | M.MagPG2 | MAG3310 | uknown | 1119 | 100% |
| | *1.462* | *M.Mag5632* | *MAGa4470* | *CG* | *285* | *96%* |
| | *1.463* | *M.Mag5632* | *MAGa4480* | *CG* | *771* | |
| **14634** | 6.630 | M.MagPG2 | MAG3310 | uknown | 1119 | 100% |
| | *6.733* | *M.MagPG2* | *MAG4250* | *CG* | *285* | *99%* |
| | *6.734* | *M.MagPG2* | *MAG4260* | *CG* | *771* | |
| **5276** | 3.355 | M.MagPG2 | MAG3310P | uknown | 1029 | 99% |
| | *1.457* | *M.Mag5632* | *MAG4470* | *CG* | *285* | *96%* |
| | *1.458* | *M.Mag5632* | *MAG4480* | *CG* | *771* | |
| **13377*** | 16.426 | M.MboH1 | ORF688P | CCTC | 1782 | 88% |
| | 9.1088 | M.Mag5632 | MAGa4470 | CG | 285 | 98% |
| | | M.Mag5632 | MAGa4480 | CG | 771 | |
| | 13.276 | M.Mag5632 | MAGa3950 | GGNCC | 1020 | 98% |
| | 8.964 | M.MboP180 | ORF173P | RCATGY | 975 | 93% |
| **4025*** | *12.329* | *M.Mag5632* | *MAGa4470* | *CG* | *285* | *96%* |
| | *12.330* | *M.Mag5632* | *MAGa4480* | *CG* | *771* | |
| | 12.274 | M.Mag5632 | MAGa3950 | GGNCC | 1020 | 98% |
| **14668*** | *1.424* | M.Mag5632 | MAGa4470 | CG | *285* | 98% |
| | | M.Mag5632 | MAGa4480 | CG | *771* | |
| | 1.356 | M.Mag5632 | MAG3950 | GGNCC | 1020 | 99% |
| **4055*** | *2.267* | M.Mag5632 | MAGa4470 | CG | *285* | 89% |
| | | M.Mag5632 | MAGa4480 | CG | *771* | |
| | *2.224* | *M.Mag5632* | *MAG3950P* | *GGNCC* | *960* | *98%* |
| | 1.713 | M.MagPG2 | MAG3310P | | 1119 | 99% |

*italic = pseudogene*

*: strains chosen for BS sequencing because they contain dcm Mtases other than the putative CpG Mtases
