## Supplementary material for "Impacts of *Mycoplasma agalactiae* restriction-modification systems on pan-epigenome dynamics and genome plasticity": TableS2.pdf

**Table S2**. Primers used for PCR and sanger sequencing

| | Name | Sequence | Tm | Mtases and homologs | Strains |
|---|---|---|---|---|---|
| Type III | **1F_typeIII*** | TACTATAAAACAGGATTATATAGAAAAAGCTAATGC | 55°C | | |
| | **1R_type III** | TCTCCCGCTTGGGTTTAAGAT | 60°C | MAGa1570 | 4025, 5632, 13377, 14668 |
| | **2R_typeIII** | ATCTTCACCAGACCCTTCATTT | 55°C | MAGa1580 | 4025, 5632, 13377 |
| | **3R_typeIII** | CCTGAACCTCCGCCAGT | 55°C | MAG1530 | 4021, 4210, 5276, 14634, PG2 |
| | **4R_typeIII** | TGGCCCTCTTTTCTTAATTCATACC | 55°C | MAG1530 | 4055 |
| | **5R_typeIII** | GCTTAATGATGCTGGAGTATATGTTCT | 55°C | MAG1580 | 14668 |
| CpG | **1F_CpG#** | AGAAGTCAATAAAGCCTATTTTACAAGGATG | 55°C | MAG4250-60 and MAGa 4470-80 | All tested strains |
| | **1R_CpG** | AAAATTCTTTCAACTTCATAAAGCAATCC | 55°C | | |

* : Universal Type III forward primer used with all Type III reverse primers and for Sanger sequencing

### : Primer used for Sanger sequencing

| PCR products |
|---|
| 655 bp (670 bp for 14668) |
| 675 bp (690 bp for 4025) |
| 670 bp |
| 675 bp (690 bp for 4025) |
| 670 bp |
| 300 bp |
