## Supplementary material for "Impacts of *Mycoplasma agalactiae* restriction-modification systems on pan-epigenome dynamics and genome plasticity": TableS3.pdf

**Table S3.** Percentage of methylated TRD couples corresponding to Type I RM systems detected b

## 5632

| | | TRD2 | | | |
|---|---|---|---|---|---|
| | | GTA | TTA | GTG | |
| TRD1 | ATC | 23% / 11% | 31% / 52% | 19% / 8% | 36% / 40% |
| | ACC | 9% / 6% | 11% / 19% | 7% / 4% | 14% / 5% |
| | | 16% / 9% | 21% / 39% | 13% / 7% | |

*5632 Type I detected methyl*

5'-ATC(N)$_5$GTA-3'

5'-ATC(N)$_5$TTA-3'

5'-ATC(N)$_5$GTG-3'

5'-ACC(N)$_5$GTA-3'

5'-ACC(N)$_5$TTA-3'

5'-ACC(N)$_5$GTG-3'

## PG2

| | | TRD2 | | |
|---|---|---|---|---|
| | | TAG | TGG | |
| TRD1 | ATC | 44% / 47% | 22% / 26% | 33% / 36% |
| | ACC | 19% / 19% | 16% / 8% | 17% / 14% |
| | | 31% / 33% | 19% / 17% | |

*PG2 Type I detected methylc*

5'-ATC(N)$_6$TAG-3'

5'-ATC(N)$_6$TGG-3'

5'-ACC(N)$_6$TAG-3'

5'-ACC(N)$_6$TGG-3'

**Abundance of detected TRD** / Abundance of predicted TRD

**number of detected TRD/total detected TRD** / number predicted TRD/total *in silico* predicted TRDs ɔy SMRT sequencing compare to *in silico* motif abundance in 5632 and PG2 stra

*ated sequences :*

*ıted sequences :*
