## Supplementary material for "Impacts of *Mycoplasma agalactiae* restriction-modification systems on pan-epigenome dynamics and genome plasticity": TableS4.pdf

**Table S4 :**Methylated motif detected in PG2 and PG2 ICEA+ variants

| | | | | | | PG2 WT | | PG2 ICEA+ | | nGenome |
|---|---|---|---|---|---|---|---|---|---|---|
| | **Detected motif** | **Consensus** | **Modification** | **Predicted motif** | **Assignated methylase** | **Fraction** | **nDetected** | **Fraction** | **nDetected** | **PG2** |
| **Type I** | AYC(N)$_6$RG | AYC(N)$_6$RG / CYA(N)$_6$GRT | m6A | - | MAG5660 | 85% | 980 | 87% | 990 | 1144 |
| | CYA(N)$_6$GRT | | | - | MAG5730 | 74% | 910 | 77% | 879 | 1144 |
| **Type II & III** | GANTC | | m6A | GANTC | MAG6680 | 86% | 4255 | 84% | 4183 | 5004 |
| | RCAC | | m6A | - | MAG1530# | 87% | 7123 | 88% | 7145 | 8081 |

### Type III Mtase
