## Supplementary material for "Impacts of *Mycoplasma agalactiae* restriction-modification systems on pan-epigenome dynamics and genome plasticity": TableS5.pdf

**Table S5** : CG abundance of M. agalactiae tested strains

| | PG2 | 5632 | 4021 | 4025 | 4055 | 4210 | 5276 | 13377 | 14634 |
|---|---|---|---|---|---|---|---|---|---|
| **Total nt** | 877439 | 944580 | 887230 | 1066370 | 1003709 | 884408 | 890157 | 1067640 | 905745 |
| Putative CpG Mtase | **-** | **+** | **-** | **+** | **-** | **-** | **-** | **+** | **-** |
| Frequence CG# | 0,01 | 0,01 | 0,01 | 0,01 | 0,01 | 0,01 | 0,01 | 0,01 | 0,01 |
| Frequence C | 0,15 | 0,15 | 0,15 | 0,15 | 0,15 | 0,15 | 0,15 | 0,15 | 0,15 |
| Frequence G | 0,14 | 0,15 | 0,14 | 0,15 | 0,14 | 0,14 | 0,14 | 0,15 | 0,14 |
| CG abundundance (Fxy)* | 0,49 | 0,46 | 0,48 | 0,49 | 0,50 | 0,49 | 0,49 | 0,49 | 0,49 |

nt = nucleotide

Mtase = Methyltransferase

### number of nucleotides or dinucleotides present in the genome /total number of nucleotides

* Based on Goto et al. (2000) = Frequence of (CG) in the genome / (Frequence of (C) * Frequence of (G))

| **14668** |
| --- |
| 967428 |
| **+** |
| 0,01 |
| 0,15 |
| 0,15 |
| 0,49 |
