## Supplementary material for "Impacts of *Mycoplasma agalactiae* restriction-modification systems on pan-epigenome dynamics and genome plasticity": TableS6.pdf

**Table S6** : Over- and under-methylated genes based on DISTAMO analysis in *M. agalactiae* 5632 and PG2 stains

### 5632 DISTAMO analysis
Overrepresented genes
Underrepresented genes

| | Genename | zScore | product | Mnemonic |
|---|---|---|---|---|
| GANTC | cdd | 4,44 | cytidine deaminase | MAGa1220 |
| | ssb | 2,75 | single-stranded DNA-binding protein | MAGa4130 |
| | vpmaE | -2,32 | variable surface lipoprotein | MAGa5850 |
| | vpmaE | -2,32 | variable surface lipoprotein | MAGa8090 |
| | vpmaW | -2,62 | vpmaW: variable surface lipoprotein | vpmaW |
| GATC | vpmaE | 3,39 | variable surface lipoprotein | MAGa5850 |
| | vpmaE | 3,39 | variable surface lipoprotein | MAGa8090 |
| | rplU | 3,17 | 50S ribosomal protein L21 | MAGa6190 |
| | MAGa4500 | 3,07 | Lipoprot | MAGa4500 |
| | MAGa2590 | 2,38 | Lipoprot | MAGa2590 |
| | MAGa3870 | 2,31 | ATP synthase C chain | MAGa3870 |
| | vpmaW | 2,11 | variable surface lipoprotein | |
| | MAGa1180 | -2 | Ribose 5 phosphate isomerase" | MAGa1180 |
| | cmk | -2,09 | Cytidylate kinase | MAGa8380 |
| | prfA | -2,19 | Peptide chain release factor 1 | MAGa8520 |
| | MAGa6690 | -2,48 | HP | MAGa6690 |
| | prs | -2,52 | ribose-phosphate pyrophosphokinase | MAGa1880 |
| GAAG | MAGa6740 | 3,15 | HP | MAGa6740 |
| | MAGa5380 | 2,91 | HP | MAGa5380 |
| | rplL | 2,77 | 50S ribosomal protein L7/L12" | rplL |
| | hit | 2,57 | HIT family protein | MAGa1990 |
| | MAGa6460 | 2,24 | GNAT family N-acetyltransferase | MAGa6460 |
| | upp | 2,19 | uracil phosphoribosyltransferase | MAGa4120 |
| | rpsI | -2,17 | 30S ribosomal protein S9 | MAGa4780 |
| | pknB | -2,19 | Serine/threonine protein kinase | MAGa2120 |
| | MAGa0310 | -2,22 | HP | MAGa0310 |
| | MAGa7610 | -2,82 | CHP | MAGa7610 |
| | MAGa1210 | -3,08 | CHP | MAGa1210 |
| GCNGC | RlmH | 3,02 | 23S rRNA (pseudouridine(1915)-N(3))-methyltransferase | MAGa1730 |
| | MAGa1720 | 2,61 | HP | MAGa1720 |
| | rplK | 2,25 | 50S ribosomal protein L11 | MaGa0860 |
| | MAga0570 | 2,08 | HP | MAga0570 |
| GGNCC | none | | | |
| AYC(N)$_5$KTR | none | | | |

### PG2 DISTAMO analysis
Overrepresented genes
Underrepresented genes

| | Genename | zScore | product | Mnemonic |
|---|---|---|---|---|
| GANTC | MAG4350 | 4,3 | Pseudogene of DNA processing protein (Smf) | MAG4350 |
| | ssb | 2,01 | single-stranded DNA-binding protein | MAG3670 |
| | infB | -2,01 | Translation initiation factor IF-2 | MAG6970 |
| | vpmaW | -3,23 | variable surface lipoprotein | |
| RCAC | MAG3140 | 4,35 | HP | MAG3140 |
| | MAG0845 | 2,96 | Hypothetical protein, truncated in C terminal | MAG0845 |
| | MAG1110 | 2,39 | Ribose 5 phosphate isomerase | MAG1110 |
| | pth | 2,31 | Peptidyl tRNA hydrolase | MAG7470 |
| | MAG6865 | 2,3 | HP | MAG6865 |
| | MAG4420 | 2,13 | HP | MAG4420 |
| | hit | 2,11 | HIT like protein (Cell cycle regulation) | MAG1990 |
| | MAG2790 | 2,08 | CHP | MAG2790 |
| | rplA | -2,09 | 50S ribosomal protein L1 | MAG0810 |
| | MAG5670 | -2,25 | CHP truncated in N term | MAG5670 |
| AYC(N)$_6$TRG | MAG4330 | -2,24 | HP lipoprotein | ompH |
