## Supplementary material for "Impacts of *Mycoplasma agalactiae* restriction-modification systems on pan-epigenome dynamics and genome plasticity": TableS7.pdf

**Table S7. Mobilome of the ten sequenced *M. agalactiae* strains**

| Strain | Mnemonic | Contig | Gene name | Function |
|---|---|---|---|---|
| **PG2** | MAG_3410 | complete genome | | Mobile element protein |
| | MAG_6150 | complete genome | | Mobile element protein |
| | MAG_7110 | complete genome | | Integrase-recombinase |
| | MAG_5690 | complete genome | | Integrase |
| | MAG6440 | complete genome | | Putative prophage protein (ps3) |
| | MAG6440b | complete genome | | Putative prophage protein (ps3) second copy |
| | MAG_2380 | complete genome | cds17 | |
| | MAG_3360 | complete genome | cds22 | |
| | MAG_3610 | complete genome | cdsB | |
| | MAG_3860 | complete genome | cds22 | |
| | MAG_3890 | complete genome | cds19 | |
| | MAG_3910 | complete genome | cds17 | |
| | MAG_3920 | complete genome | cds17 | |
| | MAG_3940 | complete genome | cds16 | |
| | MAG_3980 | complete genome | cdsC | |
| | MAG_3990 | complete genome | cds12 | |
| | MAG_4040 | complete genome | cds5 | |
| | MAG_4780 | complete genome | cds17 | |
| **4210** | peg.101 | contig_1 | | Mobile element protein |
| | peg.713 | contig_1 | | Mobile element protein |
| | peg.45 | contig_1 | | Phage integrase |
| | peg.203 | contig_1 | | Integrase-recombinase |
| | peg.631 | contig_1 | | transposase |
| | peg.632 | contig_1 | | transposase |
| | peg.133 | contig_1 | | Putative prophage protein (ps3) |
| | peg.445 | contig_1 | cds17 | ICE element |
| | peg.516 | contig_1 | cds17 | ICE element |
| | peg.625 | contig_1 | cds22 | ICE element |
| | peg.626 | contig_1 | cds22 | ICE element |
| | peg.657 | contig_1 | cdsB | ICE element |
| | peg.685 | contig_1 | cds22 | ICE element |
| | peg.686 | contig_1 | cds22 | ICE element |
| | peg.689 | contig_1 | cds19 | ICE element |
| | peg.691 | contig_1 | cds17 | ICE element |
| | peg.692 | contig_1 | cds17 | ICE element |
| | peg.693 | contig_1 | cds17 | ICE element |
| | peg.694 | contig_1 | cds16 | ICE element |
| | peg.695 | contig_1 | cds15 | ICE element |
| | peg.699 | contig_1 | cdsC | ICE element |
| | peg.700 | contig_1 | cds12 | ICE element |
| | peg.706 | contig_1 | cds5 | ICE element |
| | peg.707 | contig_1 | cds5 | ICE element |
| | peg.785 | contig_1 | cds17 | ICE element |
| **4021** | peg.1 | contig_1 | | Mobile element protein |
| | peg.2 | contig_1 | | Mobile element protein |
| | peg.283 | contig_2 | | Mobile element protein |
| | peg.534 | contig_2 | | Mobile element protein |
| | peg.140 | contig_1 | | Integrase-recombinase |
| | peg.170 | contig_1 | | Integrase-recombinase |
| | peg.340 | contig_2 | | Phage integrase |
| | peg.1035 | contig_5 | | transposase |
| | peg.1036 | contig_5 | | transposase |
| | peg.252 | contig_2 | | Putative prophage protein (ps3) |
| | peg.458 | contig_2 | cds17 | ICE element |
| | peg.542 | contig_2 | cds5 | ICE element |
| | peg.548 | contig_2 | cds12 | ICE element |
| | peg.549 | contig_2 | cdsC | ICE element |
| | peg.554 | contig_2 | cds15 | ICE element |
| | peg.555 | contig_2 | cds16 | ICE element |
| | peg.556 | contig_2 | cds17 | ICE element |
| | peg.557 | contig_2 | cds17 | ICE element |
| | peg.559 | contig_2 | cds17 | ICE element |

| | | | | |
|---|---|---|---|---|
| | peg.560 | contig_2 | cds17 | ICE element |
| | peg.562 | contig_2 | cds19 | ICE element |
| | peg.565 | contig_2 | cds22 | ICE element |
| | peg.600 | contig_2 | cdsB | ICE element |
| | peg.693 | contig_3 | cds17 | ICE element |
| | peg.1028 | contig_5 | cds22 | ICE element |
| | peg.1029 | contig_5 | cds22 | ICE element |
| | peg.1060 | contig_5 | cdsB | ICE element |
| **5276** | peg.367 | contig_1 | | Mobile element protein |
| | peg.660 | contig_1 | | Mobile element protein |
| | peg.820 | contig_3 | | Mobile element protein |
| | peg.821 | contig_3 | | Mobile element protein |
| | peg.822 | contig_3 | | Mobile element protein |
| | peg.823 | contig_3 | | Mobile element protein |
| | peg.608 | contig_1 | | Integrase |
| | peg.781 | contig_2 | | Integrase-recombinase |
| | peg.691 | contig_1 | | Putative prophage protein (ps3) |
| | peg.714 | contig_2 | | Putative prophage protein (ps3) |
| | peg.715 | contig_2 | | Putative prophage protein (ps3) |
| | peg.258 | contig_1 | cds17 | ICE element |
| | peg.361 | contig_1 | cds22 | ICE element |
| | peg.362 | contig_1 | cds22 | ICE element |
| | peg.389 | contig_1 | cdsB | ICE element |
| | peg.415 | contig_1 | cds22 | ICE element |
| | peg.418 | contig_1 | cds19 | ICE element |
| | peg.420 | contig_1 | cds17 | ICE element |
| | peg.422 | contig_1 | cds17 | ICE element |
| | peg.423 | contig_1 | cds16 | ICE element |
| | peg.424 | contig_1 | cds15 | ICE element |
| | peg.428 | contig_1 | cdsC | ICE element |
| | **peg.429** | contig_1 | cds12 | ICE element |
| | **peg.435** | contig_1 | cds5 | ICE element |
| | peg.513 | contig_1 | cds17 | ICE element |
| **14634** | peg.134 | contig_1 | | Mobile element protein |
| | peg.927 | contig_3 | | Mobile element protein |
| | peg.953 | contig_3 | | Mobile element protein |
| | peg.688 | contig_1 | | Integrase-recombinase |
| | peg.689 | contig_1 | | Integrase-recombinase |
| | peg.744 | contig_2 | | Phage integrase |
| | peg.828 | contig_3 | | Integrase-recombinase |
| | peg.220 | contig_1 | | transposase |
| | peg.893 | contig_3 | | Putative prophage protein (ps3) |
| | peg.57 | contig_1 | cds17 | ICE element |
| | peg.141 | contig_1 | cds5 | ICE element |
| | peg.147 | contig_1 | cds12 | ICE element |
| | peg.148 | contig_1 | cdsC | ICE element |
| | peg.153 | contig_1 | cds15 | ICE element |
| | peg.154 | contig_1 | cds16 | ICE element |
| | peg.155 | contig_1 | cds17 | ICE element |
| | peg.156 | contig_1 | cds17 | ICE element |
| | peg.158 | contig_1 | cds17 | ICE element |
| | peg.161 | contig_1 | cds19 | ICE element |
| | peg.164 | contig_1 | cds22 | ICE element |
| | peg.165 | contig_1 | cds22 | ICE element |
| | peg.198 | contig_1 | cdsB | ICE element |
| | peg.226 | contig_1 | cds22 | ICE element |
| | peg.351 | contig_1 | cds17 | ICE element |
| | peg.425 | contig_1 | cds17 | ICE element |
| | peg.569 | contig_1 | cdsE | ICE element |
| **5632** | MAGa1590 | complete genome | | Mobile element protein |
| | MAGa5640 | complete genome | | Mobile element protein |
| | MAGa7100 | complete genome | | Mobile element protein |
| | MAGa7110 | complete genome | | Mobile element protein |
| | MAGa7420 | complete genome | | Mobile element protein |
| | MAGa5880 | complete genome | | Integrase-recombinase |
| | MAGa6320 | complete genome | | Phage integrase |

| | | | |
|---|---|---|---|
| MAGa6690 | complete genome | | Transposase |
| MAGa7400 | complete genome | | Putative prophage protein (ps3) |
| MAGa2700 | complete genome | cdsH | ICE element |
| MAGa2980 | complete genome | cds1 | ICE element |
| MAGa2990 | complete genome | cdsA | ICE element |
| MAGa3000 | complete genome | cds12 | ICE element |
| MAGa3010 | complete genome | cds11 | ICE element |
| MAGa3020 | complete genome | cds11 | ICE element |
| MAGa3030 | complete genome | cdsB | ICE element |
| MAGa3040 | complete genome | cdsC | ICE element |
| MAGa3050 | complete genome | cdsD | ICE element |
| MAGa3060 | complete genome | cds5 | ICE element |
| MAGa3070 | complete genome | cds7 | ICE element |
| MAGa3080 | complete genome | cds13 | ICE element |
| MAGa3090 | complete genome | cds15 | ICE element |
| MAGa3100 | complete genome | cds16 | ICE element |
| MAGa3110 | complete genome | cds16 | ICE element |
| MAGa3130 | complete genome | cds17 | ICE element |
| MAGa3140 | complete genome | cds19 | ICE element |
| MAGa3150 | complete genome | cdsE | ICE element |
| MAGa3160 | complete genome | cds14 | ICE element |
| MAGa3170 | complete genome | cdsF | ICE element |
| MAGa3190 | complete genome | cdsG | ICE element |
| MAGa3200 | complete genome | cdsH | ICE element |
| MAGa3210 | complete genome | cdsG | ICE element |
| MAGa3220 | complete genome | cds22 | ICE element |
| MAGa3670 | complete genome | cds22 | ICE element |
| MAGa3680 | complete genome | cds22 | ICE element |
| MAGa4000 | complete genome | cds22 | ICE element |
| MAGa4040 | complete genome | cds5 | ICE element |
| MAGa4070 | complete genome | cdsB | ICE element |
| MAGa4850 | complete genome | cds1 | ICE element |
| MAGa4860 | complete genome | cdsA | ICE element |
| MAGa4870 | complete genome | cds12 | ICE element |
| MAGa4880 | complete genome | cds11 | ICE element |
| MAGa4890 | complete genome | cdsB | ICE element |
| MAGa4900 | complete genome | cdsC | ICE element |
| MAGa4910 | complete genome | cdsD | ICE element |
| MAGa4920 | complete genome | cds5 | ICE element |
| MAGa4930 | complete genome | cds7 | ICE element |
| MAGa4940 | complete genome | cds13 | ICE element |
| MAGa4950 | complete genome | cds15 | ICE element |
| MAGa4960 | complete genome | cds16 | ICE element |
| MAGa4980 | complete genome | cds17 | ICE element |
| MAGa4990 | complete genome | cds19 | ICE element |
| MAGa5000 | complete genome | cdsE | ICE element |
| MAGa5010 | complete genome | cds14 | ICE element |
| MAGa5020 | complete genome | cdsF | ICE element |
| MAGa5040 | complete genome | cdsG | ICE element |
| MAGa5050 | complete genome | cdsH | ICE element |
| MAGa5060 | complete genome | cds22 | ICE element |
| MAGa5250 | complete genome | cds17 | ICE element |
| MAGa6880 | complete genome | cds22 | ICE element |
| MAGa6890 | complete genome | cdsG | ICE element |
| MAGa6900 | complete genome | cdsH | ICE element |
| MAGa6910 | complete genome | cdsG | ICE element |
| MAGa6930 | complete genome | cdsF | ICE element |
| MAGa6940 | complete genome | cds14 | ICE element |
| MAGa6950 | complete genome | cdsE | ICE element |
| MAGa6960 | complete genome | cds19 | ICE element |
| MAGa6970 | complete genome | cds17 | ICE element |
| MAGa6990 | complete genome | cds16 | ICE element |
| MAGa7000 | complete genome | cds15 | ICE element |
| MAGa7010 | complete genome | cds13 | ICE element |

| | | | | |
|---|---|---|---|---|
| | MAGa7020 | complete genome | cds7 | ICE element |
| | MAGa7030 | complete genome | cds5 | ICE element |
| | MAGa7040 | complete genome | cdsD | ICE element |
| | MAGa7050 | complete genome | cdsC | ICE element |
| | MAGa7060 | complete genome | cdsB | ICE element |
| | MAGa7070 | complete genome | cds11 | ICE element |
| | MAGa7080 | complete genome | cds12 | ICE element |
| | MAGa7090 | complete genome | cdsA | ICE element |
| | MAGa7100 | complete genome | cds1 | ICE element |
| **13377** | peg.1 | contig_1 | | Mobile element protein |
| | peg.2 | contig_1 | | Mobile element protein |
| | peg.3 | contig_1 | | Mobile element protein |
| | peg.55 | contig_1 | | Mobile element protein |
| | peg.130 | contig_1 | | Mobile element protein |
| | peg.131 | contig_1 | | Mobile element protein |
| | peg.211 | contig_15 | | Mobile element protein |
| | peg.228 | contig_15 | | Mobile element protein |
| | peg.231 | contig_15 | | Mobile element protein |
| | peg.243 | contig_15 | | Mobile element protein |
| | peg.253 | contig_15 | | Mobile element protein |
| | peg.254 | contig_15 | | Mobile element protein |
| | peg.280 | contig_16 | | Mobile element protein |
| | peg.281 | contig_16 | | Mobile element protein |
| | peg.282 | contig_16 | | Mobile element protein |
| | peg.290 | contig_16 | | Mobile element protein |
| | peg.326 | contig_17 | | Mobile element protein |
| | peg.343 | contig_17 | | Mobile element protein |
| | peg.485 | contig_5 | | Mobile element protein |
| | peg.626 | contig_7 | | Mobile element protein |
| | peg.645 | contig_7 | | Mobile element protein |
| | peg.676 | contig_7 | | Mobile element protein |
| | peg.712 | contig_8 | | Mobile element protein |
| | peg.795 | contig_8 | | Mobile element protein |
| | peg.848 | contig_8 | | Mobile element protein |
| | peg.863 | contig_8 | | Mobile element protein |
| | peg.878 | contig_8 | | Mobile element protein |
| | peg.879 | contig_8 | | Mobile element protein |
| | peg.1054 | contig_8 | | Mobile element protein |
| | peg.891 | contig_8 | | Integrase-recombinase |
| | peg.223 | contig_15 | | transposase |
| | peg.63 | contig_1 | | Putative prophage protein (ps3) |
| | peg.255 | contig_15 | | Phage helicase |
| | peg.262 | contig_15 | | phage DNA polymerase |
| | peg.264 | contig_15 | | DNA primase, phage associated |
| | peg.265 | contig_15 | | Phage DNA primase |
| | peg.696 | contig_7 | | phage DNA polymerase |
| | peg.697 | contig_7 | | DNA primase, phage associated |
| | peg.4 | contig_1 | cds22 | ICE element |
| | peg.5 | contig_1 | cds7 | ICE element |
| | peg.6 | contig_1 | cds5 | ICE element |
| | peg.7 | contig_1 | cds5 | ICE element |
| | peg.8 | contig_1 | cds5 | ICE element |
| | peg.9 | contig_1 | cdsD | ICE element |
| | peg.10 | contig_1 | cdsD | ICE element |
| | peg.12 | contig_1 | cdsC | ICE element |
| | peg.13 | contig_1 | cdsC | ICE element |
| | peg.14 | contig_1 | cdsB | ICE element |
| | peg.15 | contig_1 | cds11 | ICE element |
| | peg.16 | contig_1 | cds11 | ICE element |
| | peg.17 | contig_1 | cds12 | ICE element |
| | peg.18 | contig_1 | cdsA | ICE element |
| | peg.20 | contig_1 | cds1 | ICE element |
| | peg.21 | contig_1 | cds1 | ICE element |
| | peg.205 | contig_15 | cds22 | ICE element |
| | peg.206 | contig_15 | cds22 | ICE element |
| | peg.271 | contig_16 | cdsA | ICE element |

| | | | | |
|---|---|---|---|---|
| | peg.272 | contig_16 | cds12 | ICE element |
| | peg.273 | contig_16 | cds11 | ICE element |
| | peg.274 | contig_16 | cdsB | ICE element |
| | peg.275 | contig_16 | cdsC | ICE element |
| | peg.276 | contig_16 | cdsD | ICE element |
| | peg.277 | contig_16 | cds5 | ICE element |
| | peg.278 | contig_16 | cds7 | ICE element |
| | peg.279 | contig_16 | cds22 | ICE element |
| | peg.320 | contig_17 | cds5 | ICE element |
| | peg.321 | contig_17 | cds5 | ICE element |
| | peg.327 | contig_17 | cdsB | ICE element |
| | peg.345 | contig_18 | cds19 | ICE element |
| | peg.346 | contig_18 | cds19 | ICE element |
| | peg.347 | contig_18 | cds17 | ICE element |
| | peg.348 | contig_18 | cds17 | ICE element |
| | peg.349 | contig_18 | cds16 | ICE element |
| | peg.350 | contig_18 | cds17 | ICE element |
| | peg.352 | contig_18 | cds19 | ICE element |
| | peg.354 | contig_18 | cds22 | ICE element |
| | peg.371 | contig_20 | cdsH | ICE element |
| | peg.402 | contig_5 | cds1 | ICE element |
| | peg.404 | contig_5 | cdsA | ICE element |
| | peg.405 | contig_5 | cds12 | ICE element |
| | peg.406 | contig_5 | cds11 | ICE element |
| | peg.407 | contig_5 | cdsB | ICE element |
| | peg.408 | contig_5 | cdsC | ICE element |
| | peg.409 | contig_5 | cdsD | ICE element |
| | peg.410 | contig_5 | cds5 | ICE element |
| | peg.411 | contig_5 | cds7 | ICE element |
| | peg.413 | contig_5 | cds15 | ICE element |
| | peg.414 | contig_5 | cds16 | ICE element |
| | peg.415 | contig_5 | cds22 | ICE element |
| | peg.492 | contig_5 | cdsF | ICE element |
| | peg.494 | contig_5 | cdsG | ICE element |
| | peg.495 | contig_5 | cdsG | ICE element |
| | peg.496 | contig_5 | cdsH | ICE element |
| | peg.497 | contig_5 | cdsG | ICE element |
| | peg.498 | contig_5 | cds22 | ICE element |
| | peg.507 | contig_5 | cdsH | ICE element |
| | peg.585 | contig_5 | cds17 | ICE element |
| | peg.592 | contig_5 | cds1 | ICE element |
| | peg.594 | contig_5 | cds5 | ICE element |
| | peg.597 | contig_5 | cdsC | ICE element |
| | peg.599 | contig_5 | cds12 | ICE element |
| | peg.603 | contig_5 | cds15 | ICE element |
| | peg.604 | contig_5 | cds22 | ICE element |
| | peg.605 | contig_5 | cdsG | ICE element |
| | peg.606 | contig_5 | cdsH | ICE element |
| | peg.607 | contig_5 | cdsG | ICE element |
| | peg.609 | contig_5 | cdsF | ICE element |
| | peg.613 | contig_5 | cds14 | ICE element |
| | peg.614 | contig_5 | cdsE | ICE element |
| | peg.615 | contig_5 | cds19 | ICE element |
| | peg.616 | contig_5 | cds17 | ICE element |
| | peg.618 | contig_5 | cds16 | ICE element |
| | peg.790 | contig_8 | cds17 | ICE element |
| **4025** | peg.171 | contig_1 | | Mobile element protein |
| | peg.172 | contig_1 | | Mobile element protein |
| | peg.194 | contig_12 | | Mobile element protein |
| | peg.197 | contig_12 | | Mobile element protein |
| | peg.198 | contig_12 | | Mobile element protein |
| | peg.199 | contig_12 | | Mobile element protein |
| | peg.254 | contig_12 | | Mobile element protein |
| | peg.255 | contig_12 | | Mobile element protein |
| | peg.259 | contig_12 | | Mobile element protein |
| | peg.263 | contig_12 | | Mobile element protein |
| | peg.311 | contig_12 | | Mobile element protein |

| | | | |
|---|---|---|---|
| peg.312 | contig_12 | | Mobile element protein |
| peg.315 | contig_12 | | Mobile element protein |
| peg.323 | contig_12 | | Mobile element protein |
| peg.363 | contig_12 | | Mobile element protein |
| peg.388 | contig_12 | | Mobile element protein |
| peg.396 | contig_12 | | Mobile element protein |
| peg.400 | contig_12 | | Mobile element protein |
| peg.401 | contig_12 | | Mobile element protein |
| peg.402 | contig_12 | | Mobile element protein |
| peg.445 | contig_12 | | Mobile element protein |
| peg.446 | contig_12 | | Mobile element protein |
| peg.448 | contig_12 | | Mobile element protein |
| peg.488 | contig_12 | | Mobile element protein |
| peg.506 | contig_12 | | Mobile element protein |
| peg.507 | contig_12 | | Mobile element protein |
| peg.509 | contig_12 | | Mobile element protein |
| peg.547 | contig_12 | | Mobile element protein |
| peg.565 | contig_12 | | Mobile element protein |
| peg.573 | contig_12 | | Mobile element protein |
| peg.581 | contig_12 | | Mobile element protein |
| peg.616 | contig_12 | | Mobile element protein |
| peg.622 | contig_12 | | Mobile element protein |
| peg.623 | contig_12 | | Mobile element protein |
| peg.645 | contig_12 | | Mobile element protein |
| peg.742 | contig_14 | | Mobile element protein |
| peg.752 | contig_2 | | Mobile element protein |
| peg.796 | contig_2 | | Mobile element protein |
| peg.798 | contig_2 | | Mobile element protein |
| peg.799 | contig_2 | | Mobile element protein |
| peg.800 | contig_2 | | Mobile element protein |
| peg.803 | contig_2 | | Mobile element protein |
| peg.816 | contig_2 | | Mobile element protein |
| peg.900 | contig_4 | | Mobile element protein |
| peg.931 | contig_4 | | Mobile element protein |
| peg.932 | contig_4 | | Mobile element protein |
| peg.933 | contig_4 | | Mobile element protein |
| peg.494 | contig_12 | | Integrase |
| peg.705 | contig_14 | | Integrase-recombinase |
| peg.535 | contig_12 | | Transposase |
| peg.603 | contig_12 | | Putative prophage protein (ps3) |
| peg.153 | contig_1 | cds1 | ICE element |
| peg.155 | contig_1 | cds5 | ICE element |
| peg.159 | contig_1 | cds12 | ICE element |
| peg.161 | contig_1 | cds12 | ICE element |
| peg.164 | contig_1 | cds5 | ICE element |
| peg.166 | contig_1 | cds1 | ICE element |
| peg.278 | contig_12 | cds22 | ICE element |
| peg.283 | contig_12 | cds5 | ICE element |
| peg.287 | contig_12 | cdsB | ICE element |
| peg.387 | contig_12 | cds17 | ICE element |
| **peg.543** | contig_12 | cds22 | ICE element |
| **peg.548** | contig_12 | cds19 | ICE element |
| **peg.550** | contig_12 | cds17 | ICE element |
| **peg.551** | contig_12 | cds16 | ICE element |
| **peg.552** | contig_12 | cds15 | ICE element |
| **peg.556** | contig_12 | cds12 | ICE element |
| **peg.557** | contig_12 | cdsC | ICE element |
| **peg.558** | contig_12 | cds12 | ICE element |
| **peg.561** | contig_12 | cds5 | ICE element |
| **peg.563** | contig_12 | cds1 | ICE element |
| peg.583 | contig_12 | cdsH | ICE element |
| peg.584 | contig_12 | cdsH | ICE element |
| peg.749 | contig_2 | cds19 | ICE element |
| peg.750 | contig_2 | cds19 | ICE element |
| peg.756 | contig_2 | cds22 | ICE element |
| peg.786 | contig_2 | cdsH | ICE element |
| peg.863 | contig_2 | cds17 | ICE element |

| | | | | |
|---|---|---|---|---|
| | peg.878 | contig_2 | cdsH | ICE element |
| | peg.907 | contig_4 | cds1 | ICE element |
| | peg.911 | contig_4 | cds5 | ICE element |
| | peg.912 | contig_4 | cds5 | ICE element |
| | peg.913 | contig_4 | cds5 | ICE element |
| | peg.920 | contig_4 | cds5 | ICE element |
| | peg.921 | contig_4 | cds5 | ICE element |
| | peg.925 | contig_4 | cds1 | ICE element |
| | peg.926 | contig_4 | cds1 | ICE element |
| | peg.958 | contig_8 | cds1 | ICE element |
| | peg.963 | contig_8 | cds5 | ICE element |
| | peg.964 | contig_8 | cds5 | ICE element |
| | peg.970 | contig_8 | cds5 | ICE element |
| | peg.971 | contig_8 | cds5 | ICE element |
| | peg.972 | contig_8 | cds5 | ICE element |
| | peg.973 | contig_8 | cds5 | ICE element |
| | peg.974 | contig_8 | cds5 | ICE element |
| | peg.980 | contig_8 | cds1 | ICE element |
| | peg.981 | contig_8 | cds1 | ICE element |
| **14668** | peg.258 | contig_1 | | Mobile element protein |
| | peg.259 | contig_1 | | Mobile element protein |
| | peg.433 | contig_1 | | Mobile element protein |
| | peg.614 | contig_1 | | Mobile element protein |
| | peg.615 | contig_1 | | Mobile element protein |
| | peg.792 | contig_2 | | Mobile element protein |
| | peg.657 | contig_1 | | integrase-recombinase protein |
| | peg.685 | contig_1 | | Integrase-recombinase |
| | peg.626 | contig_1 | | DNA helicase, phage-associated |
| | peg.636 | contig_1 | | DNA primase, phage associated # P4-type |
| | peg.650 | contig_1 | | Phage tail fiber protein |
| | peg.653 | contig_1 | | Phage portal protein |
| | peg.654 | contig_1 | | Phage terminase, large subunit |
| | peg.768 | contig_2 | | Putative prophage protein (ps3) |
| | peg.242 | contig_1 | cds17 | ICE element |
| | peg.330 | contig_1 | cds22 | ICE element |
| | peg.331 | contig_1 | cds22 | ICE element |
| | peg.361 | contig_1 | cds22 | ICE element |
| | peg.365 | contig_1 | cds19 | ICE element |
| | peg.367 | contig_1 | cds17 | ICE element |
| | peg.368 | contig_1 | cds16 | ICE element |
| | peg.369 | contig_1 | cds15 | ICE element |
| | peg.373 | contig_1 | cdsC | ICE element |
| | peg.374 | contig_1 | cdsC | ICE element |
| | peg.375 | contig_1 | cds12 | ICE element |
| | peg.377 | contig_1 | cdsA | ICE element |
| | peg.380 | contig_1 | cds5 | ICE element |
| | peg.383 | contig_1 | cdsB | ICE element |
| | peg.479 | contig_1 | cds17 | ICE element |
| | peg.789 | contig_2 | cdsH | ICE element |
| | peg.790 | contig_2 | cdsH | ICE element |
| **4055** | peg.60 | contig_1 | | Mobile element protein |
| | peg.61 | contig_1 | | Mobile element protein |
| | peg.64 | contig_1 | | Mobile element protein |
| | peg.65 | contig_1 | | Mobile element protein |
| | peg.68 | contig_1 | | Mobile element protein |
| | peg.178 | contig_1 | | Mobile element protein |
| | peg.403 | contig_1 | | Mobile element protein |
| | peg.405 | contig_1 | | Mobile element protein |
| | peg.699 | contig_1 | | Mobile element protein |
| | peg.702 | contig_1 | | Mobile element protein |
| | peg.708 | contig_1 | | Mobile element protein |
| | peg.816 | contig_1 | | Mobile element protein |
| | peg.817 | contig_1 | | Mobile element protein |
| | peg.869 | contig_1 | | Mobile element protein |
| | peg.874 | contig_1 | | Mobile element protein |
| | peg.931 | contig_1 | | Mobile element protein |
| | peg.980 | contig_1 | | Mobile element protein |

| | | | | |
|---|---|---|---|---|
| | peg.983 | contig_1 | | Mobile element protein |
| | peg.987 | contig_1 | | Mobile element protein |
| | peg.1028 | contig_1 | | Mobile element protein |
| | peg.1032 | contig_1 | | Mobile element protein |
| | peg.1130 | contig_1 | | Mobile element protein |
| | peg.1131 | contig_1 | | Mobile element protein |
| | peg.1133 | contig_1 | | Mobile element protein |
| | peg.1322 | contig_1 | | Mobile element protein |
| | peg.1323 | contig_1 | | Mobile element protein |
| | peg.1334 | contig_1 | | Mobile element protein |
| | peg.1394 | contig_1 | | Mobile element protein |
| | peg.1396 | contig_1 | | Mobile element protein |
| | peg.1533 | contig_1 | | Mobile element protein |
| | peg.1535 | contig_1 | | Mobile element protein |
| | peg.1548 | contig_1 | | Mobile element protein |
| | peg.1570 | contig_1 | | Mobile element protein |
| | peg.1608 | contig_1 | | Mobile element protein |
| | peg.1626 | contig_1 | | Mobile element protein |
| | peg.1646 | contig_1 | | Mobile element protein |
| | peg.1671 | contig_1 | | Mobile element protein |
| | peg.1740 | contig_1 | | Mobile element protein |
| | peg.1748 | contig_1 | | Mobile element protein |
| | peg.1750 | contig_1 | | Mobile element protein |
| | peg.1772 | contig_1 | | Mobile element protein |
| | peg.1782 | contig_1 | | Mobile element protein |
| | peg.1866 | contig_1 | | Mobile element protein |
| | peg.1891 | contig_1 | | Mobile element protein |
| | peg.1895 | contig_1 | | Mobile element protein |
| | peg.1952 | contig_1 | | Mobile element protein |
| | peg.1956 | contig_1 | | Mobile element protein |
| | peg.1959 | contig_1 | | Mobile element protein |
| | peg.1963 | contig_1 | | Mobile element protein |
| | peg.1964 | contig_1 | | Mobile element protein |
| | peg.1966 | contig_1 | | Mobile element protein |
| | peg.1969 | contig_1 | | Mobile element protein |
| | peg.1975 | contig_1 | | Mobile element protein |
| | peg.1976 | contig_1 | | Mobile element protein |
| | peg.1978 | contig_1 | | Mobile element protein |
| | peg.2352 | contig_1 | | Mobile element protein |
| | peg.2551 | contig_1 | | Mobile element protein |
| | peg.2731 | contig_1 | | Mobile element protein |
| | peg.2732 | contig_1 | | Mobile element protein |
| | peg.2735 | contig_1 | | Mobile element protein |
| | peg.2749 | contig_1 | | Mobile element protein |
| | peg.2808 | contig_1 | | Mobile element protein |
| | peg.2810 | contig_1 | | Mobile element protein |
| | peg.2866 | contig_1 | | Mobile element protein |
| | peg.2869 | contig_1 | | Mobile element protein |
| | peg.2905 | contig_1 | | Mobile element protein |
| | peg.1791 | contig_1 | | integrase-recombinase protein |
| | peg.2399 | contig_1 | | Transposase ISMmy1I |
| | peg.497 | contig_1 | | Putative prophage protein (ps3) |
| | peg.498 | contig_1 | | Putative prophage protein (ps3) |
| | peg.496 | contig_1 | cds14 | ICE element |
| | peg.1418 | contig_1 | cds5 | ICE element |
