## Supplementary material for "Impacts of *Mycoplasma agalactiae* restriction-modification systems on pan-epigenome dynamics and genome plasticity": TableS8.pdf

**Table S8.** BLASTP results for M. agalactiae active methyltransferases against bacteria other than the Mollicutes class

| Methyltransferases * | Description | Scientific Name | Max Score | Total Score | Query Cover | E value | Per. Ident | Acc. Len | Accession |
|---|---|---|---|---|---|---|---|---|---|
| MAGa2700 | Dam family site-specific DNA-(adenine-N6)-methyltransferase [Solobacterium sp.] | Solobacterium sp. | 338 | 338 | 96% | 1,00E-113 | 59.34% | 273 | MBF1103133.1 |
| | Dam family site-specific DNA-(adenine-N6)-methyltransferase [Solobacterium sp.] | Solobacterium sp. | 336 | 336 | 96% | 5,00E-113 | 59.34% | 273 | MBF1078290.1 |
| | Dam family site-specific DNA-(adenine-N6)-methyltransferase [Solobacterium sp.] | Solobacterium sp. | 335 | 335 | 96% | 1,00E-112 | 58.97% | 273 | MBF1086133.1 |
| | DNA adenine methylase [Macrococcus caseolyticus] | Macrococcus caseolyticus | 332 | 332 | 96% | 2,00E-111 | 59.11% | 269 | RKO15992.1 |
| | DNA adenine methylase [Macrococcus sp. IME1552] | Macrococcus sp. IME1552 | 331 | 331 | 96% | 8,00E-111 | 59.11% | 269 | WP_096077300.1 |
| | DNA adenine methylase [Macrococcus caseolyticus] | Macrococcus caseolyticus | 325 | 650 | 96% | 1,00E-108 | 58.36% | 269 | WP_099483891.1 |
| | Dam family site-specific DNA-(adenine-N6)-methyltransferase [Macrococcus canis] | Macrococcus canis | 325 | 325 | 96% | 2,00E-108 | 57.99% | 269 | WP_164941655.1 |
| | DNA adenine methylase [Megamonas hypermegale] | Megamonas hypermegale | 324 | 324 | 96% | 4,00E-108 | 58.30% | 276 | WP_027889300.1 |
| | Modification methylase DpnIIA [Megamonas hypermegale] | Megamonas hypermegale | 324 | 324 | 96% | 4,00E-108 | 58.30% | 278 | SNU95784.1 |
| | MULTISPECIES: Dam family site-specific DNA-(adenine-N6)-methyltransferase [Staphylococcus] | Staphylococcus | 323 | 323 | 96% | 7,00E-108 | 56.88% | 269 | WP_071560813.1 |
| MAGa7650 | site-specific DNA-methyltransferase [Clostridia bacterium] | Clostridia bacterium | 469 | 469 | 99% | 3,00E-162 | 61.44% | 376 | NLO90117.1 |
| | site-specific DNA-methyltransferase [Treponema vincentii] | Treponema vincentii | 463 | 463 | 98% | 6,00E-160 | 61.02% | 372 | WP_006187946.1 |
| | site-specific DNA-methyltransferase [Treponema vincentii] | Treponema vincentii | 462 | 462 | 98% | 1,00E-159 | 61.29% | 372 | WP_016518474.1 |
| | site-specific DNA-methyltransferase [Campylobacter curvus] | Campylobacter curvus | 459 | 459 | 96% | 2,00E-158 | 62.47% | 371 | WP_169783906.1 |
| | site-specific DNA-methyltransferase [Clostridium isatidis] | Clostridium isatidis | 458 | 458 | 93% | 4,00E-158 | 62.61% | 359 | WP_119866576.1 |
| | site-specific DNA-methyltransferase [Campylobacter concisus] | Campylobacter concisus | 455 | 455 | 97% | 1,00E-156 | 60.60% | 369 | WP_107832996.1 |
| | MULTISPECIES: site-specific DNA-methyltransferase [Campylobacter] | Campylobacter | 454 | 454 | 97% | 2,00E-156 | 60.87% | 369 | WP_009294648.1 |
| | MULTISPECIES: site-specific DNA-methyltransferase [unclassified Campylobacter] | unclassified Campylobacter | 454 | 454 | 97% | 2,00E-156 | 59.67% | 369 | WP_086237322.1 |
| | site-specific DNA-methyltransferase [Marinitoga sp. 1138] | Marinitoga sp. 1138 | 453 | 453 | 97% | 4,00E-156 | 60.48% | 372 | WP_175418024.1 |
| | site-specific DNA-methyltransferase [Campylobacter hyointestinalis] | Campylobacter hyointestinalis | 452 | 452 | 97% | 1,00E-155 | 59.40% | 370 | WP_147499997.1 |
| MAGa4250 | DNA cytosine methyltransferase [Fusobacterium sp. CM21] | Fusobacterium sp. CM21 | 469 | 469 | 100% | 1,00E-163 | 70.79% | 351 | WP_032841141.1 |
| | M.Fnu4HI [Fusobacterium nucleatum] | Fusobacterium nucleatum | 469 | 469 | 100% | 1,00E-163 | 70.48% | 351 | ADX97301.1 |
| | DNA (cytosine-5-)-methyltransferase [Parvimonas sp. S3374] | Parvimonas sp. S3374 | 458 | 458 | 99% | 2,00E-159 | 71.34% | 347 | WP_201275014.1 |
| | DNA cytosine methyltransferase [Parvimonas sp. S3374] | Parvimonas sp. S3374 | 458 | 458 | 99% | 3,00E-159 | 71.34% | 346 | MBK1467862.1 |
| | DNA cytosine methyltransferase [Bacillus toyonensis] | Bacillus toyonensis | 456 | 456 | 99% | 3,00E-159 | 67.52% | 315 | WP_098944381.1 |
| | DNA cytosine methyltransferase [Sphingobium sp. TCM1] | Sphingobium sp. TCM1 | 454 | 454 | 99% | 3,00E-158 | 67.52% | 315 | WP_066863026.1 |
| | DNA cytosine methyltransferase [Bacillus mycoides] | Bacillus mycoides | 452 | 452 | 99% | 2,00E-157 | 67.52% | 315 | WP_016127586.1 |
| | RecName: Full=Modification methylase Bsp6I; Short=M.Bsp6I; AltName: Full=Cytosine-specific methyltransfe | Bacillus sp. RFL6 | 450 | 450 | 99% | 9,00E-157 | 66.56% | 315 | P43420.1 |
| | DNA (cytosine-5-)-methyltransferase [Hazenella sp. IB182357] | Hazenella sp. IB182357 | 448 | 448 | 99% | 1,00E-155 | 67.20% | 324 | WP_191142604.1 |
| | TPA: DNA cytosine methyltransferase [Bacilli bacterium] | Bacilli bacterium | 441 | 441 | 99% | 1,00E-152 | 64.76% | 332 | HHU19175.1 |
| MAGa3950 | DNA (cytosine-5-)-methyltransferase [Epulopiscium sp. Nele67-Bin005] | Epulopiscium sp. Nele67-Bin005 | 510 | 510 | 98% | 1,00E-179 | 70.57% | 345 | OON95436.1 |
| | DNA (cytosine-5-)-methyltransferase [Bisgaard taxon 44 str. B96_4] | Bisgaard taxon 44 str. B96_4 | 501 | 501 | 96% | 3,00E-176 | 73.62% | 341 | RIY34047.1 |
| | DNA (cytosine-5-)-methyltransferase [Campylobacter sp. RM15925] | Campylobacter sp. RM15925 | 499 | 499 | 98% | 4,00E-175 | 73.05% | 343 | WP_169941814.1 |
| | DNA (cytosine-5-)-methyltransferase [Fusobacterium periodonticum 2_1_31] | Fusobacterium periodonticum 2_ | 495 | 495 | 97% | 2,00E-173 | 70.18% | 361 | KGE62123.1 |
| | DNA cytosine methyltransferase [Fusobacterium nucleatum] | Fusobacterium nucleatum | 494 | 494 | 97% | 3,00E-173 | 70.48% | 344 | WP_098702880.1 |
| | DNA (cytosine-5-)-methyltransferase [Fusobacterium periodonticum D10] | Fusobacterium periodonticum D1 | 494 | 494 | 97% | 4,00E-173 | 70.18% | 361 | EKA93595.1 |
| | Eco47II family restriction endonuclease [Pseudoleptotrichia goodfellowii] | Pseudoleptotrichia goodfellowii | 502 | 502 | 97% | 1,00E-172 | 72.29% | 600 | MBF4804993.1 |
| | DNA cytosine methyltransferase [Campylobacter jejuni] | Campylobacter jejuni | 490 | 490 | 98% | 7,00E-172 | 70.96% | 338 | EAI4070807.1 |
| | DNA methyltransferase [Campylobacter jejuni] | Campylobacter jejuni | 490 | 490 | 98% | 1,00E-171 | 70.96% | 347 | AXL47160.1 |
| | DNA cytosine methyltransferase [Campylobacter jejuni] | Campylobacter jejuni | 489 | 489 | 98% | 3,00E-171 | 70.66% | 338 | WP_002921457.1 |
| 13377_426 | DNA (cytosine-5-)-methyltransferase [[Micrococcus] candicans] | [Micrococcus] candicans | 822 | 822 | 99% | 0.0 | 65.43% | 594 | WP_198687347.1 |
| | DNA (cytosine-5-)-methyltransferase [Carnobacterium alterfunditum] | Carnobacterium alterfunditum | 805 | 805 | 99% | 0.0 | 64.02% | 600 | WP_081884459.1 |
| | DNA (cytosine-5-)-methyltransferase [Staphylococcus sp. HMSC34C02] | Staphylococcus sp. HMSC34C02 | 796 | 796 | 99% | 0.0 | 63.74% | 599 | WP_070854712.1 |
| | DNA-methyltransferase Dcm [Mycobacteroides abscessus subsp. abscessus] | Mycobacteroides abscessus subs | 795 | 795 | 99% | 0.0 | 64.47% | 599 | SIH36365.1 |
| | DNA (cytosine-5-)-methyltransferase [Staphylococcus haemolyticus] | Staphylococcus haemolyticus | 793 | 793 | 99% | 0.0 | 63.56% | 599 | WP_080402264.1 |
| | MULTISPECIES: DNA (cytosine-5-)-methyltransferase [Staphylococcus] | Staphylococcus | 793 | 793 | 99% | 0.0 | 63.56% | 599 | WP_070822985.1 |
| | DNA (cytosine-5-)-methyltransferase [Staphylococcus haemolyticus] | Staphylococcus haemolyticus | 792 | 792 | 99% | 0.0 | 63.39% | 599 | WP_085060987.1 |
| | DNA (cytosine-5-)-methyltransferase [Staphylococcus haemolyticus] | Staphylococcus haemolyticus | 792 | 792 | 99% | 0.0 | 63.39% | 599 | WP_117287751.1 |
| | DNA (cytosine-5-)-methyltransferase [Staphylococcus haemolyticus] | Staphylococcus haemolyticus | 790 | 790 | 99% | 0.0 | 63.39% | 599 | WP_080367091.1 |
| | DNA (cytosine-5-)-methyltransferase [Staphylococcus epidermidis] | Staphylococcus epidermidis | 789 | 789 | 99% | 0.0 | 63.45% | 598 | WP_115339039.1 |
| 4025_583 | GCATC--recognizing Type II restriction modification system (MmyCIII) adenine DNA methyltransferase subun | synthetic Mycoplasma mycoides . | 439 | 439 | 97% | 1,00E-150 | 61.94% | 362 | ADH21784.1 |
| | modification methylase [Lactococcus garvieae] | Lactococcus garvieae | 382 | 382 | 91% | 6,00E-128 | 56.25% | 398 | PCS02374.1 |
| | Dam family site-specific DNA-(adenine-N6)-methyltransferase [Streptococcus uberis] | Streptococcus uberis | 391 | 391 | 92% | 3,00E-127 | 59.23% | 709 | WP_154631601.1 |
| | Dam family site-specific DNA-(adenine-N6)-methyltransferase [Streptococcus parauberis] | Streptococcus parauberis | 385 | 385 | 92% | 5,00E-125 | 57.44% | 711 | WP_139058238.1 |
| | D12 class N6 adenine-specific DNA methyltransferase family protein [Streptococcus pneumoniae NP070] | Streptococcus pneumoniae NP07 | 375 | 375 | 91% | 6,00E-125 | 56.89% | 405 | EHD56120.1 |
| | D12 class N6 adenine-specific DNA methyltransferase family protein [Streptococcus pneumoniae GA44128] | Streptococcus pneumoniae GA44 | 375 | 375 | 0,91 | 7E-125 | 56.89% | 406 | EHZ51383.1 |
| | DNA adenine methylase [Streptococcus parauberis NCFD 2020] | Streptococcus parauberis NCFD | 385 | 385 | 0,92 | 8E-125 | 57.44% | 720 | EGE54547.1 |
| | Dam family site-specific DNA-(adenine-N6)-methyltransferase [Vagococcus penaei] | Vagococcus penaei | 383 | 383 | 0,9 | 6E-124 | 56.63% | 708 | WP_126844542.1 |
| | DNA adenine methylase [Streptococcus pneumoniae] | Streptococcus pneumoniae | 376 | 376 | 0,91 | 9E-124 | 57.19% | 509 | WP_057607080.1 |

| | | | | | | | | | |
|---|---|---|---|---|---|---|---|---|---|
| | Adenine-specific DNA methylase [Enterococcus durans] | Enterococcus durans | 372 | 372 | 0,94 | 1E-123 | 56.65% | 398 | STQ48463.1 |

BLASTP on NCBI (https://blast.ncbi.nlm.nih.gov)
Eclude Mollicutes option
Cut-off E-value of <0.001
Filter : Identity between 100% and 50% / Coverage between 100% and 50%

* : CpG methylase, Type III related MTases (MAGa1570, MAGa1580 and MAG1530) and the Type II methylase MTase 13377_964
are not represented in this table as they have no homologs outside Mollicutes class
