## Supplementary figures and images for "Impacts of *Mycoplasma agalactiae* restriction-modification systems on pan-epigenome dynamics and genome plasticity"

### FigureS1.tif

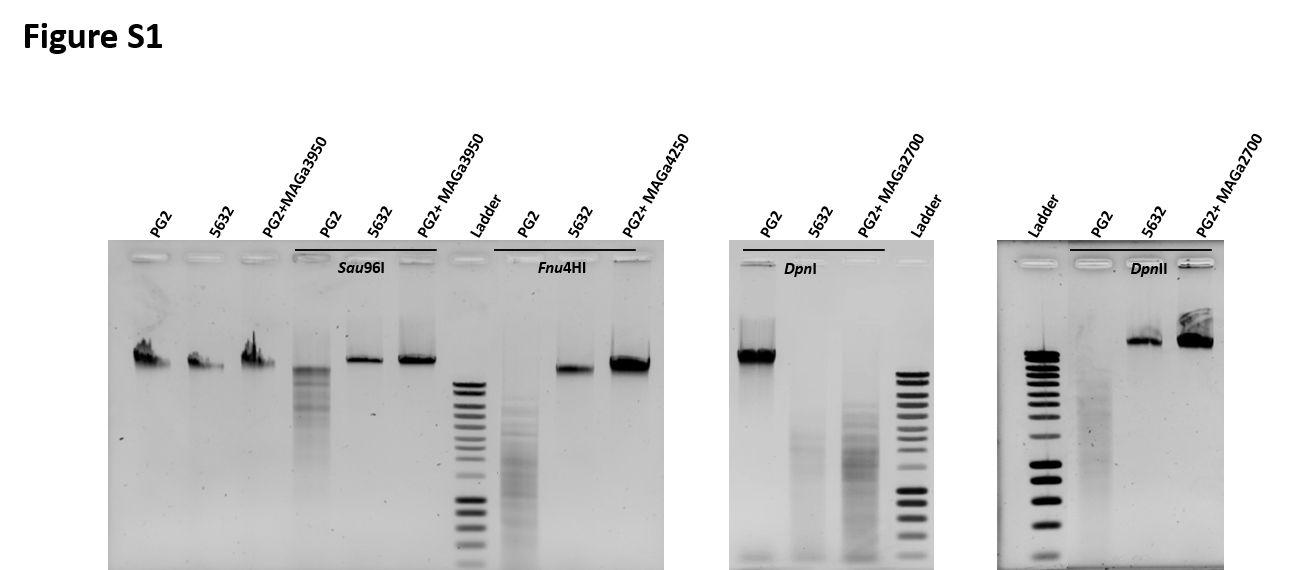

### FigureS2.tif

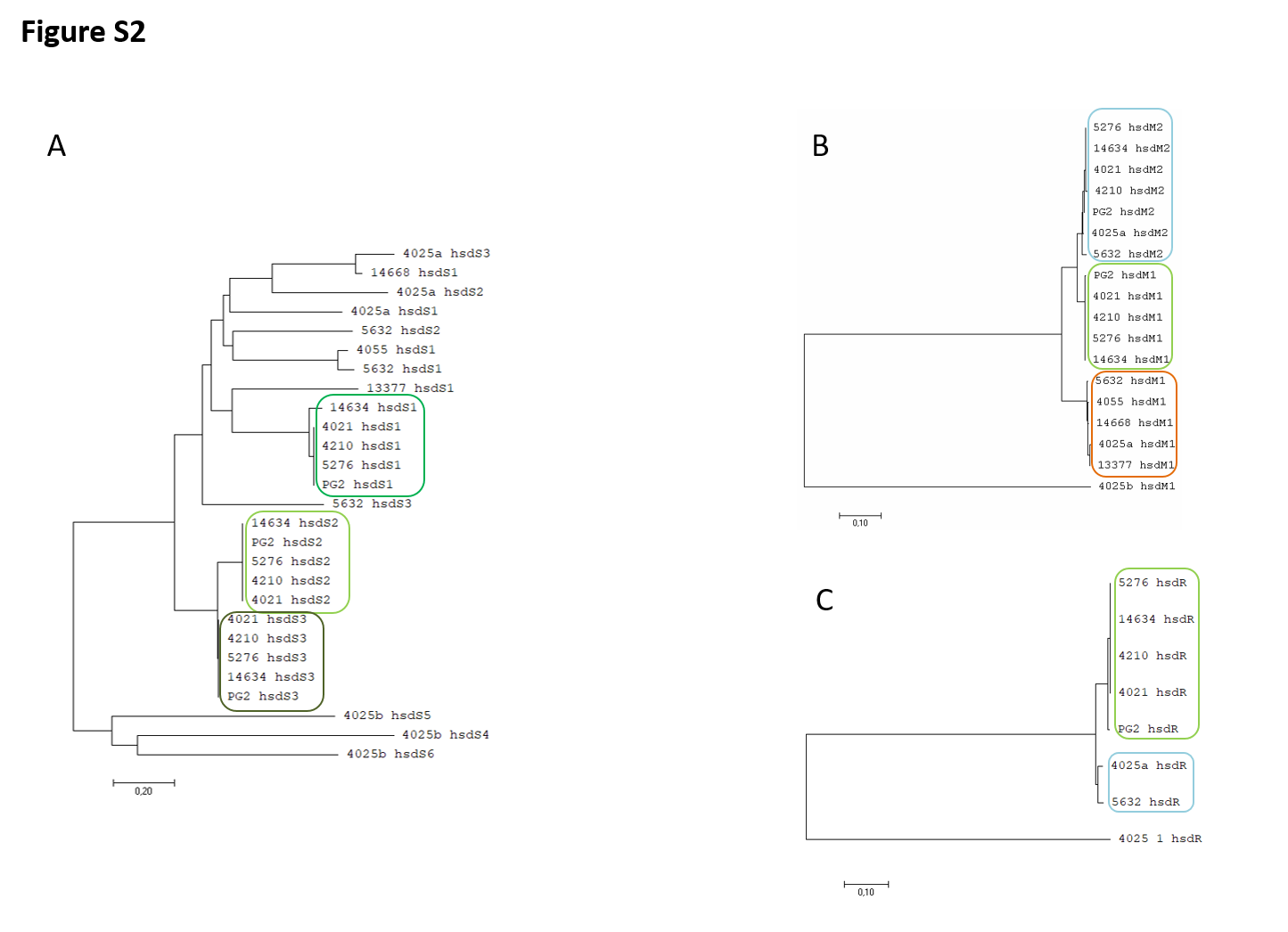

### FigureS3.tif

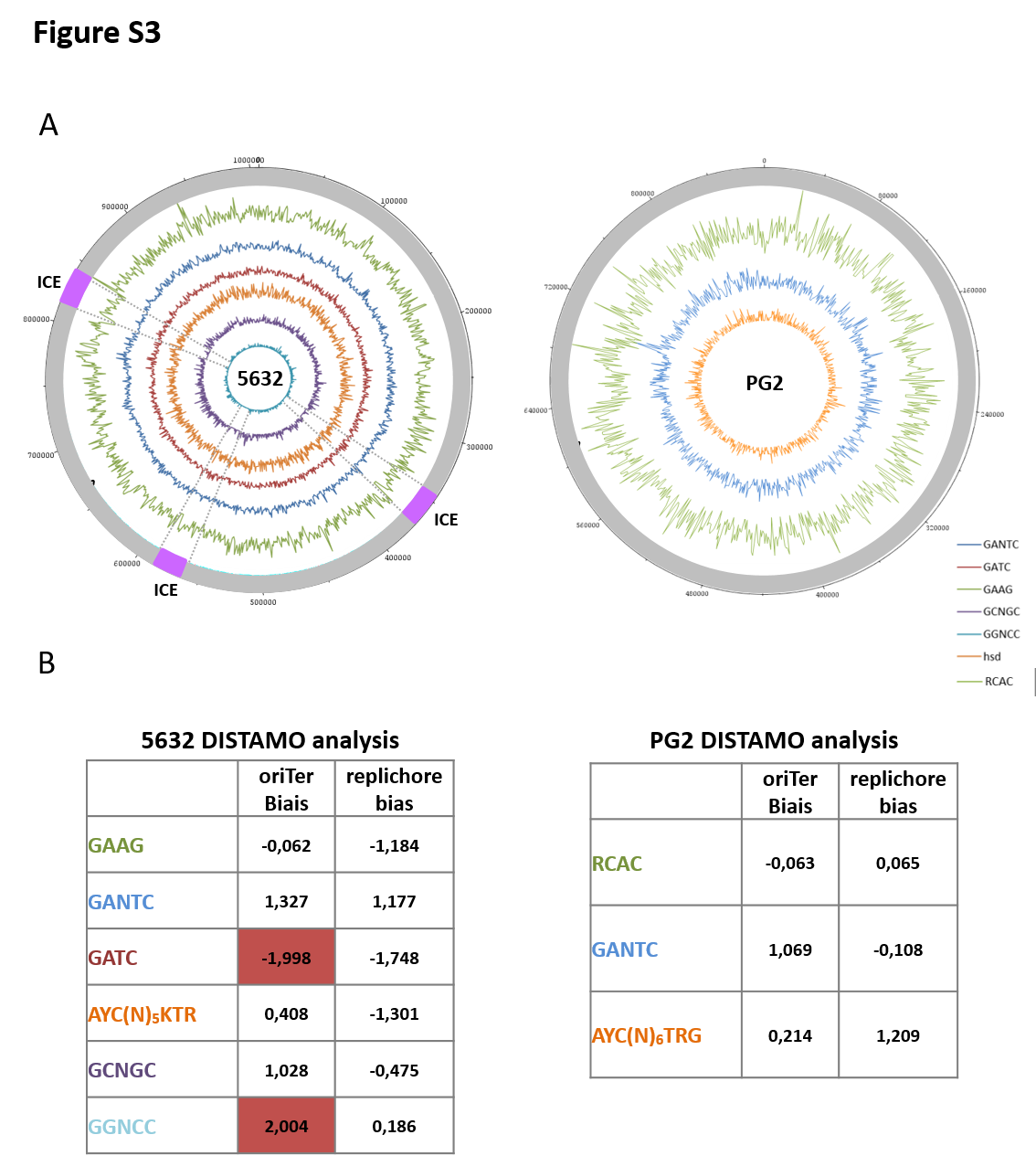
